# Influenza A virus initiates genome selection through promoter-dependent nuclear export

**DOI:** 10.64898/2026.08.14.744863

**Authors:** Kuang-Yu Chen, Alison Rep, Loïc Carrique, Fangzheng Wang, Ecco Staller, Jonathan M. Grimes, Ervin Fodor

## Abstract

Influenza A virus replicates its segmented RNA genome in the nucleus in the context of viral ribonucleoprotein (vRNP) complexes. Newly synthesised vRNPs are exported to the cytoplasm for virion assembly, whereas structurally similar complementary RNPs (cRNPs), which function as replication intermediates, are not incorporated into virions. The molecular basis for this selectivity remains unclear. Here, combining promoter mutagenesis, single-molecule fluorescence in situ hybridisation (smFISH), virus-like particle (VLP) assays and cryo-electron microscopy (cryo-EM), we find that cRNPs are retained in the nucleus and that export competence is encoded by promoter architecture, with the length of the single-stranded 3′ promoter region as the principal contributing feature and promoter sequence and duplex architecture further modulating export efficiency. cRNPs carrying export-competent promoters are incorporated into virus-like particles, demonstrating a functional link between nuclear export and packaging competence. Cryo-EM analyses reveal that both 3′ vRNA and cRNA promoters bind a common polymerase surface site but are associated with distinct polymerase conformational landscapes, with only the vRNA promoter being compatible with an encapsidase conformation of the polymerase. Together, our findings identify a promoter-dependent checkpoint that links RNA promoter architecture and polymerase conformation to RNP export and packaging. These results demonstrate that influenza virus initiates genome selection in the nucleus and reveal nuclear export as an early step in genome selection.

## Introduction

The packaging of segmented RNA genomes presents a fundamental challenge for RNA viruses, which must selectively assemble complete sets of genome segments into progeny virions. To ensure genome integrity, many segmented RNA viruses rely on specific RNA signals and ribonucleoprotein architectures that discriminate genome segments from replication intermediates or aberrant RNA species during assembly. A particular challenge arises when genomic RNAs coexist with structurally similar replication intermediates that must be excluded from progeny virions. How viruses achieve this selectivity remains an outstanding question in RNA virus biology.

Influenza A virus provides a well-studied model for selective genome packaging. Its genome comprises eight negative-sense single-stranded viral RNA (vRNA) segments, each encapsidated by multiple copies of nucleoprotein (NP) and a single heterotrimeric RNA-dependent RNA polymerase complex composed of PB2, PB1 and PA, together forming a viral ribonucleoprotein complex (vRNP)^1^. During infection, vRNPs serve as templates for transcription to generate capped viral mRNAs and for replication to produce complementary RNA (cRNA) intermediates^2–4^. Similar to vRNA, cRNA is encapsidated by the viral polymerase complex and NP to form complementary ribonucleoprotein complexes (cRNPs), which act as templates for vRNA synthesis during the second step of genome replication. Despite their structural and compositional similarity to vRNPs^5^, cRNPs are not incorporated into virions^6,7^, indicating that influenza virus possesses a mechanism capable of distinguishing genomic RNPs from replication intermediates.

While biochemical and genetic studies established that influenza virus selectively packages vRNA, the molecular basis for this selectivity remains unresolved. Previous work focused largely on cytoplasmic events, including selective segment-segment interactions that contribute to assembly of the characteristic 7+1 genome bundle^8,9^. However, influenza virus is unusual among RNA viruses in that genome replication occurs in the host cell nucleus, raising the possibility that genome selection begins before RNPs reach the cytoplasm.

Following transcription and replication, newly synthesised vRNPs are exported from the nucleus to the cytoplasm for virion assembly. This process is mediated by the viral nuclear export protein (NEP, also known as NS2), which is essential for vRNP nuclear export^10^. NEP contains leucine-rich nuclear export signals that engage the cellular export receptor CRM1/XPO1, thereby enabling vRNPs to transit through the nuclear pore complex^11,12^. Early models proposed a “daisy-chain” mechanism in which the viral matrix protein M1 bridges NEP and vRNPs via interactions with NP^13,14^. Subsequent studies demonstrated that NEP can also associate directly with vRNPs, through interactions with the viral polymerase, while still engaging M1, suggesting that multiple, cooperative binding mechanisms may contribute to vRNP export^15^. More recently, structural and biochemical analyses identified a direct interaction between NEP and the influenza virus polymerase at the interface of the PA C-terminal domain (PA-C) and PB1^16,17^. While NEP binding at this interface modulates viral RNA synthesis, it remains unclear how this interaction supports nuclear export.

Each influenza A virus vRNA segment contains untranslated regions (UTRs) at both termini comprising highly conserved promoter sequences of thirteen nucleotides at the 5′ end and twelve nucleotides at the 3′ end, followed by segment-specific sequences. Partial complementarity between the 5′ and 3′ termini enables the viral promoter to adopt defined secondary structures that are essential for transcription and replication. Extensive mutational analyses have shown that disruption of promoter base-pairing or alteration of promoter length profoundly affect RNA synthesis, underscoring the functional importance of promoter architecture^18–23^.

Recent advances in cryo-electron microscopy (cryo-EM) have provided detailed structural insights into promoter-bound influenza polymerase complexes^24–28^. These studies revealed that the first ten nucleotides of the 5′ vRNA promoter form a conserved stem-loop (“hook”) structure that binds within a pocket at the PA-PB1 interface of the polymerase. The distal portion of the 5′ vRNA promoter, beginning at position 11, base-pairs with the 3′ vRNA promoter to form a short duplex. In contrast, the first nine nucleotides of the 3′ vRNA promoter remain single-stranded and adopt distinct conformations depending on the functional state of the polymerase. During transcription and replication initiation, the 3′ end is positioned in the template entry channel towards the polymerase active site, whereas in alternative configurations it binds to distinct sites, mode A or mode B site, on the surface of the polymerase^29^.

Structures of cRNA promoter-bound polymerase complexes reveal a broadly similar overall organisation, with the 5′ cRNA hook occupying the same PA-PB1 pocket as vRNA. However, the 3′ cRNA promoter differs in both length and behaviour: it contains a longer single-stranded region and has been observed either bound at the mode B site on the surface or positioned in a pre-catalytic state within the template entry channel^24,29^. Together, these observations support a model in which the 5′ promoter adopts a stable stem-loop conformation, while the 3′ promoter contains a flexible single-stranded region, nine nucleotides in vRNA and eleven nucleotides in cRNA, that enables dynamic transitions between surface-bound and active-site-engaged states during transcription and genome replication^24,27,28^.

Here, we investigate how influenza virus discriminates between vRNPs and cRNPs during nuclear export. Using single-molecule fluorescence in situ hybridisation (smFISH) in a reconstituted RNP system, we directly visualise viral RNA localisation. By combining promoter mutagenesis, functional virus-like particle (VLP) assays, and cryo-EM analysis of NEP-bound polymerase complexes, we identify promoter architecture as the key determinant of export competence. These results establish a direct link between promoter architecture, nuclear export and genome packaging, providing a mechanistic framework for how influenza A virus distinguishes genomic vRNPs from replication intermediates.

## Results

### cRNPs are retained in the nucleus

Influenza A virus selectively packages vRNPs into progeny virions while excluding cRNPs, which serve as intermediates during genome replication. To determine whether cRNPs are exported from the nucleus, we established an RNP reconstitution system combined with single-molecule fluorescence in situ hybridisation (smFISH) to visualise viral RNAs. In this system, we co-express the three subunits of the influenza virus RNA polymerase, PB1, PB2, and PA, together with NP and segment six vRNA, to reconstitute vRNPs. To ensure that all positive-sense RNA signal detected by smFISH could be attributed exclusively to cRNA rather than viral mRNA, which shares the same polarity, we used an endonuclease-inactive PA mutant (PA D108A)^30^ to abolish cap-dependent transcription. Primer extension analysis confirmed the absence of mRNA under these conditions (Fig. 1a, b).

**Figure 1.**
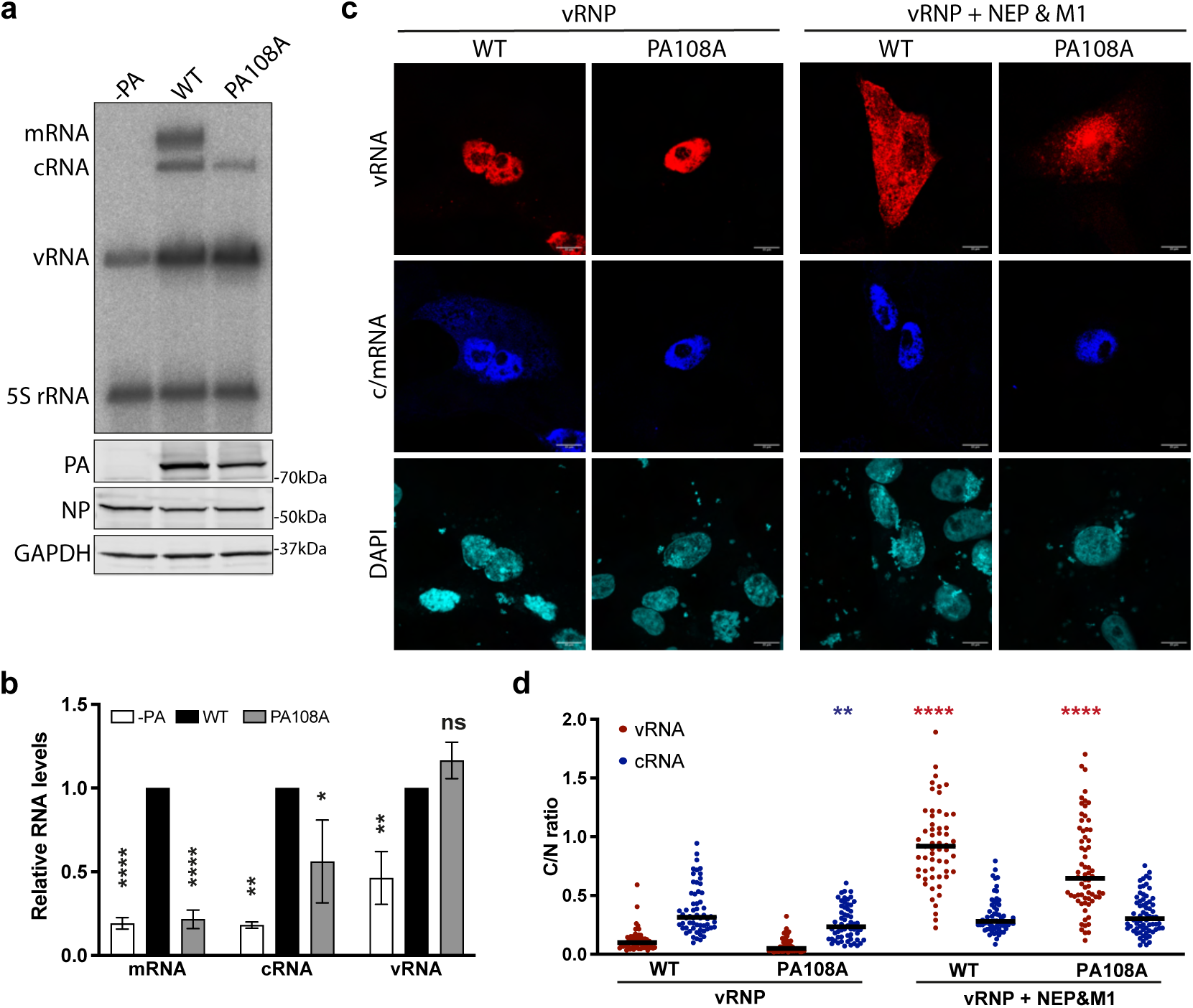
vRNA, but not cRNA, is exported from the nucleus. **a** Primer extension analysis of viral RNAs from RNP reconstitution experiments. HEK-293T cells were transfected with plasmids expressing PB2, PB1, PA (WT or PA D108A), NP, and a vRNA template, and harvested 24 h post-transfection for RNA isolation. PA was omitted as a negative control. Western blot analysis was performed to assess protein expression using antibodies against PA, NP, and GAPDH. **b** Quantification of mRNA, cRNA, and vRNA levels from the primer extension analysis shown in **a**. RNA levels were normalised to WT condition. Error bars represent the mean ± SD (n = 3). Statistical analysis was performed using one-way ANOVA with multiple comparisons. **c** Representative smFISH images of Vero E6 cells expressing RNP components. RNA localisation in cells expressing PA D108A was compared with that in cells expressing WT PA in the presence or absence of NEP and M1. smFISH was performed using probes specific for vRNA (red, upper panels) and cRNA/mRNA (blue, middle panels). Nuclear DNA was stained with DAPI (lower panels). **d** Quantification of the cytoplasmic-to-nuclear (C/N) mean fluorescence intensity ratio from individual cells across two biological replicates: WT (n = 61), PA D108A (n = 59), WT (+NEP&M1) (n = 61), and PA D108A (+NEP&M1) (n = 66). Black lines indicate the mean. Statistical analysis was performed using one-way ANOVA with multiple comparisons relative to the corresponding WT vRNA or cRNA value.

We next examined the intracellular localisation of viral RNAs in the RNP reconstitution assay by using smFISH. In the absence of the viral export factor NEP and M1, vRNA was confined to the nucleus in both the wild-type and PA D108A reconstitution systems (Fig. 1c, d). Positive-sense RNA generated by the wild-type polymerase was detected in both the nucleus and cytoplasm, consistent with the presence of exported viral mRNA. In contrast, positive-sense RNA generated by the PA D108A mutant was restricted to the nucleus, indicating that cRNA is not exported.

To determine whether cRNPs can be exported in the presence of the viral export machinery, we co-expressed NEP and M1. Under these conditions, vRNA accumulated in both the nucleus and cytoplasm, consistent with export of vRNPs. In contrast, positive-sense RNA remained confined to the nucleus in the PA D108A background despite expression of NEP and M1 (Fig. 1c, d). Quantification of cytoplasmic-to-nuclear fluorescence intensity ratios confirmed efficient export of vRNA but not cRNA (Fig. 1d).

These findings demonstrate that cRNPs are selectively retained in the nucleus even under conditions that support vRNP export. Thus, discrimination between genomic and complementary RNPs occurs prior to virion assembly and suggests that nuclear export represents an early checkpoint in influenza virus genome selection.

### Promoter RNA determines RNP nuclear export

Because vRNPs and cRNPs contain the same protein components and differ only in the sequence of their associated RNA, we next asked whether the terminal promoter architecture determines export competence. To test this, we generated a chimeric RNA template in which both the 5′ and 3′ promoter regions of the vRNA were replaced with the corresponding cRNA promoter sequences (Fig. 2a, mutant 1).

**Figure 2.**
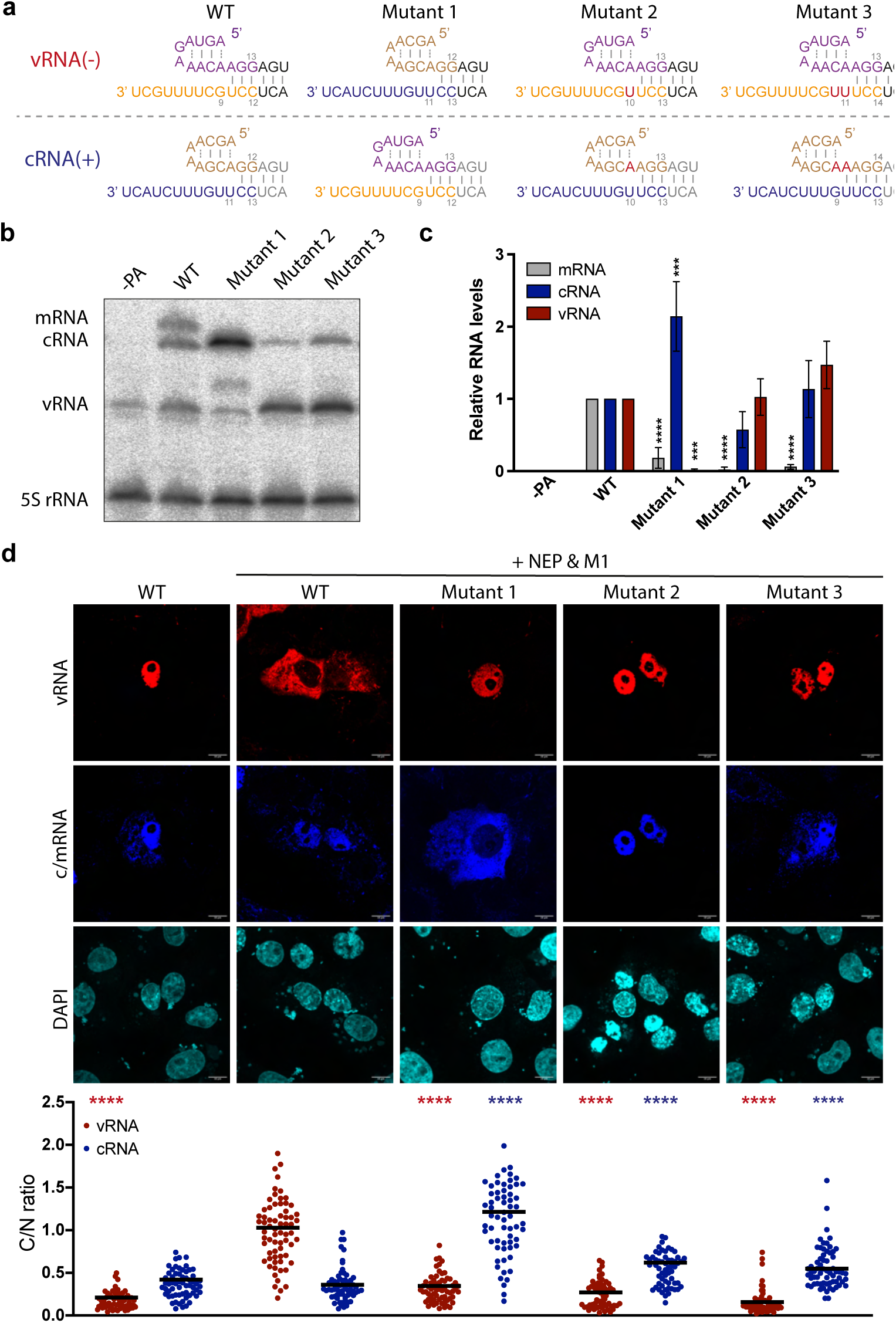
Promoter identity determines RNP nuclear export. **a** Schematic representation of the wild-type and mutant vRNA and cRNA promoter sequences. Sequences are color-coded as follows: 5′ vRNA positions 1-13 (purple), 3′ vRNA positions 1-12 (orange), 5′ cRNA positions 1-13 (brown), 3′ cRNA positions 1-12 (blue), vRNA promoter duplex (black), cRNA promoter duplex (grey), and inserted nucleotides (red). **b** Primer extension analysis of viral RNAs from RNP reconstitution experiments. HEK-293T cells were transfected with plasmids expressing PB2, PB1, PA, NP, and wild-type or mutant NA vRNA templates (WT and mutants 1-3). Cells were harvested 24 h post-transfection for RNA isolation. PA was omitted as a negative control. **c** Quantification of mRNA, cRNA, and vRNA levels from the primer extension analysis shown in **b**. RNA levels were normalised to those in the WT condition. Error bars represent the mean ± SD (n = 3). Statistical analysis was performed using one-way ANOVA with multiple comparisons relative to WT. **d** Representative smFISH images of Vero E6 cells transfected for RNP reconstitution in the presence of NEP and M1. smFISH was performed to detect vRNA (red) and cRNA (blue) localisation. Nuclei were stained with DAPI. Quantification of the cytoplasmic-to-nuclear (C/N) mean fluorescence intensity ratio from individual cells across two biological replicates: WT (n = 61), WT (+NEP&M1) (n = 67), mutant 1 (n = 63), mutant 2 (n = 59), mutant 3 (n = 65). Black lines indicate the mean. Statistical analysis was performed using one-way ANOVA with multiple comparisons relative to the WT vRNA or cRNA value.

RNA synthesis was first assessed by primer extension analysis in the absence of NEP and M1 to determine the intrinsic effects of promoter replacement on polymerase activity. Promoter exchange abolished detectable mRNA synthesis, increased cRNA accumulation and reduced vRNA levels (Fig. 2b, c). Similar effects were observed in the presence of NEP and M1, under conditions corresponding to those used in the smFISH experiments (Supplementary Fig. 1).

We next examined RNA localisation by smFISH. Strikingly, promoter exchange reversed the export phenotype: whereas wild-type vRNA accumulated in the cytoplasm following expression of NEP and M1, promoter-swapped vRNA remained confined to the nucleus, while cRNA accumulated in the cytoplasm (Fig. 2d). Thus, promoter exchange reversed the export behaviour of both RNA species.

The vRNA and cRNA promoters differ not only in sequence but also in the length of the single-stranded region at the 3′ end upstream of the promoter duplex. This region comprises nine nucleotides in vRNA and eleven nucleotides in cRNA. We therefore asked whether promoter length contributes to export competence. To test this, we inserted one or two uridine residues at position 9 of the vRNA 3′ promoter (Fig. 2a, mutants 2 and 3), extending the single-stranded region from nine to ten or eleven nucleotides, respectively. These changes simultaneously shortened the corresponding single-stranded region of the cRNA 3′ promoter from eleven to ten or nine nucleotides.

Primer extension analysis performed in the absence of NEP and M1 showed that both mutants remained replication competent but failed to produce detectable mRNA (Fig. 2b, c). Comparable effects were observed in the presence of NEP and M1 (Supplementary Fig. 1). smFISH analysis revealed that extension of the vRNA 3′ promoter abolished vRNP export. cRNPs remained predominantly nuclear, although both mutants exhibited increased cytoplasmic-to-nuclear ratios of cRNA relative to wild-type (Fig. 2d).

Together, these results identify promoter architecture as a key determinant of RNP nuclear export, demonstrating that export competence follows promoter identity rather than RNA polarity. A negative-sense vRNA carrying cRNA promoter sequences was retained in the nucleus, whereas a positive-sense cRNA carrying vRNA promoter sequences became export competent. The length of the single-stranded 3′ promoter region contributes substantially to this discrimination, as extension of the vRNA promoter abolished export and shortening of the cRNA promoter partially relieved nuclear retention. However, shortening the cRNA 3′ promoter alone was insufficient to confer robust export competence, indicating that additional promoter features contribute to the differential export of vRNPs and cRNPs.

### The 3′ promoter sequence contributes to vRNP nuclear export

Having established that promoter identity determines RNP export competence, we next asked whether specific sequence differences between the vRNA and cRNA promoters contribute to this process. The vRNA and cRNA promoters differ at several positions within the single-stranded 3′ promoter region. To assess the contribution of these differences, we introduced mutations at positions 3 and 8 (mutant 4) or at positions 3, 5 and 8 (mutant 5) into the vRNA 3′ promoter. Corresponding substitutions were introduced into the cRNA 3′ promoter (mutants 6 and mutant 7) (Fig. 3a). In addition, reciprocal mutations were introduced simultaneously into both promoters at positions 3 and 8 (mutant 8) or positions 3, 5 and 8 (mutant 9) (Fig. 4a). Positions 3 and 8 were mutated together to preserve the complementarity required for formation of the corresponding 5′ RNA hook structure^20,26^.

**Figure 3.**
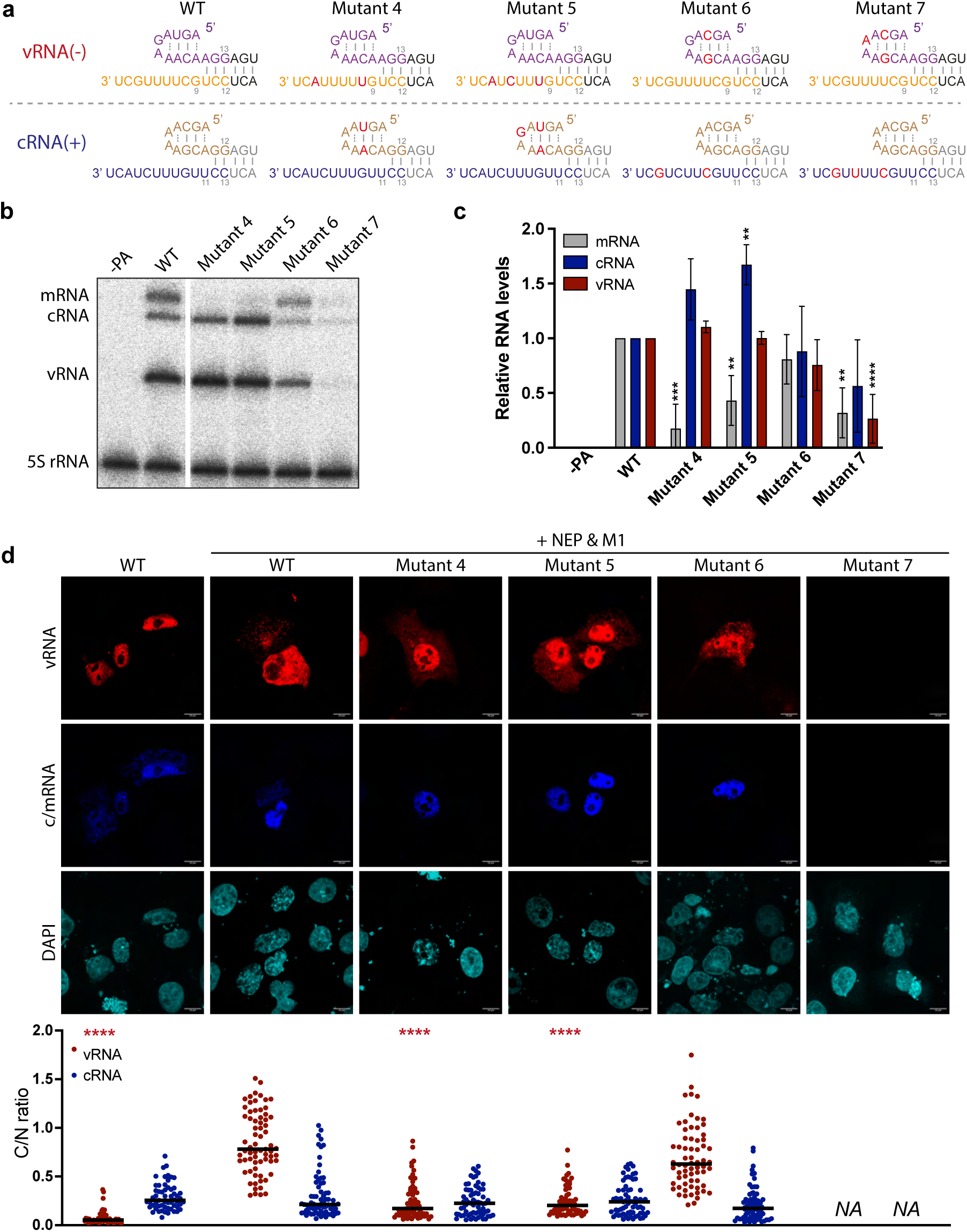
Mutations at positions 3, 5, and 8 at 3′ vRNA promoter or 3′ cRNA promoter affect nuclear export. **a** Schematic representation of the wild-type and mutant vRNA and cRNA promoter sequences. Sequences are colour-coded as in Fig. 2A, and mutations are highlighted in red. **b** Primer extension analysis of viral RNAs from RNP reconstitution experiments using wild-type and mutant NA vRNA templates (WT and mutants 4-7). **C** Quantification of mRNA, cRNA, and vRNA levels from **b**. **d** Representative smFISH images and quantification of the cytoplasmic-to-nuclear (C/N) mean fluorescence intensity ratio from individual cells across two biological replicates: WT (n = 57), WT (+NEP&M1) (n = 69), mutant 4 (n = 67), mutant 5 (n = 67), and mutant 6 (n = 66). Experimental procedures and statistical analyses were performed as described in Fig. 2.

**Figure 4.**
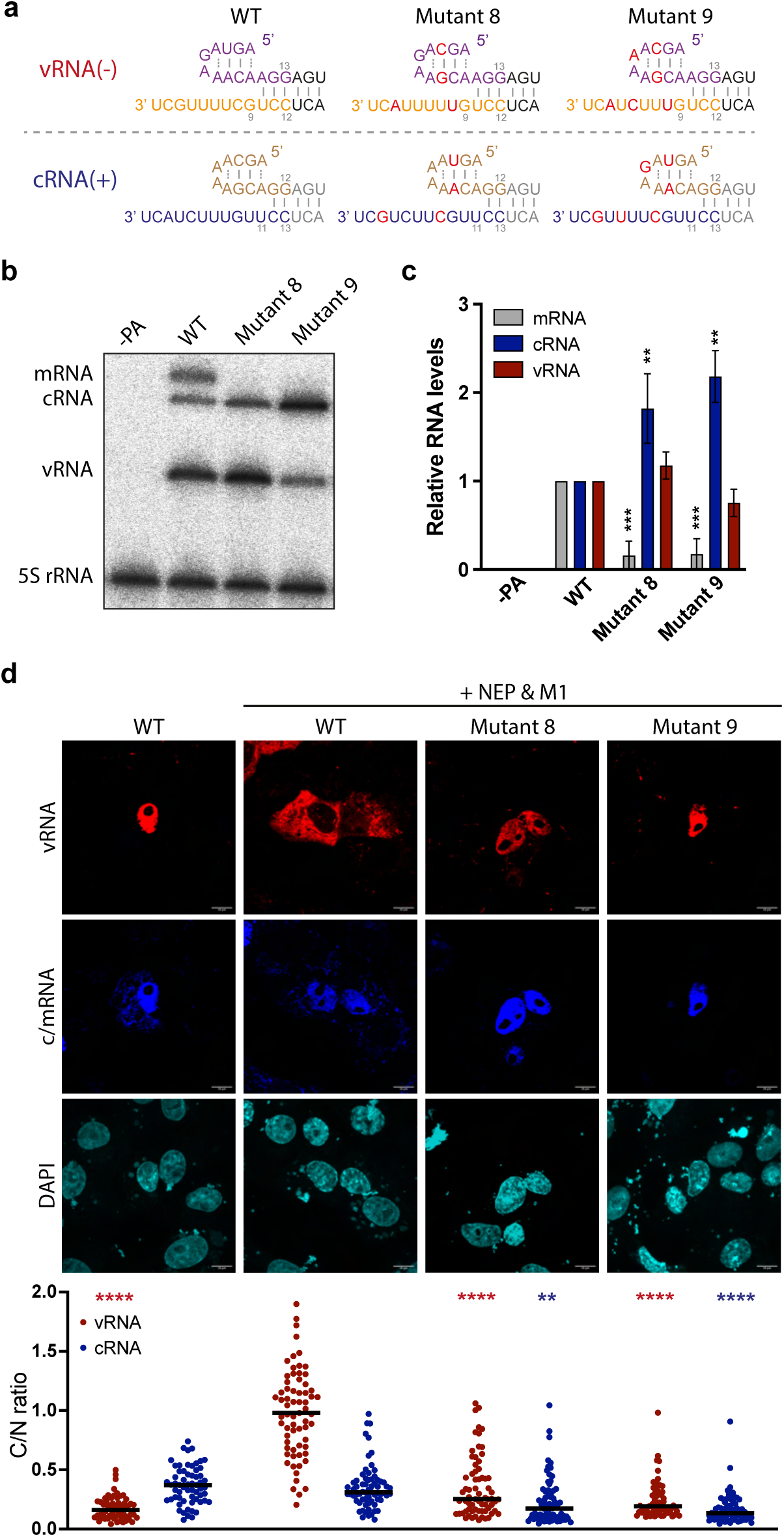
Reciprocal mutations introduced simultaneously at positions 3, 5, and 8 in both 3′ vRNA promoter or 3′ cRNA promoter disrupt nuclear export. **a** Schematic representation of the wild-type and mutant vRNA and cRNA promoter sequences. Sequences are colour-coded as in Fig. 2a, and mutations are highlighted in red. **b** Primer extension analysis of viral RNAs from RNP reconstitution experiments using wild-type and mutant NA vRNA templates (WT and mutants 8 and 9). **c** Quantification of mRNA, cRNA, and vRNA levels from (B). **d** Representative smFISH images and quantification of the cytoplasmic-to-nuclear (C/N) mean fluorescence intensity ratio from individual cells across two biological replicates: WT (n = 61), WT (+NEP&M1) (n = 67), mutant 8 (n = 68), and mutant 9 (n = 65). Experimental procedures and statistical analyses were performed as described in Fig. 2.

RNA synthesis was first assessed by primer extension analysis in the absence of NEP and M1. All mutant templates, with the exception of mutant 7, remained replication competent. In contrast, transcription was substantially reduced for all mutants except mutant 6 (Fig. 3b, c and Fig. 4b, c). Similar effects were observed in the presence of NEP and M1 (Supplementary Fig. 2).

We next examined intracellular localisation of the mutant RNAs by smFISH. Mutants 4 and 5, carrying cRNA-specific substitutions within the vRNA 3′ promoter, showed significantly reduced but detectable accumulation of vRNA in the cytoplasm relative to wild-type, while cRNA remained confined to the nucleus (Fig. 3d). Mutant 6, carrying substitutions within the cRNA 3′ promoter, exhibited a localisation pattern indistinguishable from wild-type. Mutant 7 produced reduced RNA levels insufficient for reliable quantification. Mutants 8 and 9, in which reciprocal substitutions were introduced into both the vRNA and cRNA promoters, showed significant reduction in cytoplasmic vRNA although mutant 8 exhibited detectable levels. Both mutants also showed significantly reduced cRNA accumulation possibly due to some residual mRNA in the wild-type contributing to the positive sense-specific signal (Fig. 4d).

Taken together, these findings indicate that the sequence of the 3′ vRNA promoter contributes to efficient vRNP export, although export activity is still partially retained when it is altered. Mutations at positions 3 and 5 of the 5′ vRNA promoter do not significantly affect vRNP export, indicating these residues are dispensable for export. The 5′ vRNA promoter is primarily required for polymerase activity rather than export, as substitution with the cRNA hook abolishes polymerase function. Overall, these findings indicate that promoter sequence modulates nuclear export competence but is not sufficient to determine it.

### A nine-nucleotide 3′ promoter region confers export competence

Our experiments above demonstrated that both the length and sequence of the 3′ promoter influence nuclear export. To investigate the role of promoter length further, we generated mutants in which the nine-nucleotide single-stranded region of the vRNA 3′ promoter was replaced with either ten (mutant 10) or eleven (mutant 11) nucleotides from the cRNA 3′ promoter. These substitutions simultaneously converted the corresponding 5′ cRNA promoter into a vRNA-like promoter and shortened the single-stranded region at the 3′ end of cRNA to ten or nine nucleotides, respectively (Fig. 5a). Conversely, we replaced the eleven-nucleotide single-stranded region of the cRNA 3′ promoter with either ten (mutant 12) or nine (mutant 13) nucleotides from the vRNA 3′ promoter. These substitutions converted the corresponding 5′ vRNA promoter into a cRNA-like promoter and increased the length of the single-stranded region at the 3′ end of vRNA to ten or eleven nucleotides, respectively (Fig. 5a). To preserve the conserved adenine at position 10 of the vRNA 5′ promoter, which is required for formation of the 5′ hook structure^20,26^, the complementary nucleotide at position 10 of the cRNA 3′ promoter was changed from cytosine to uracil.

**Figure 5.**
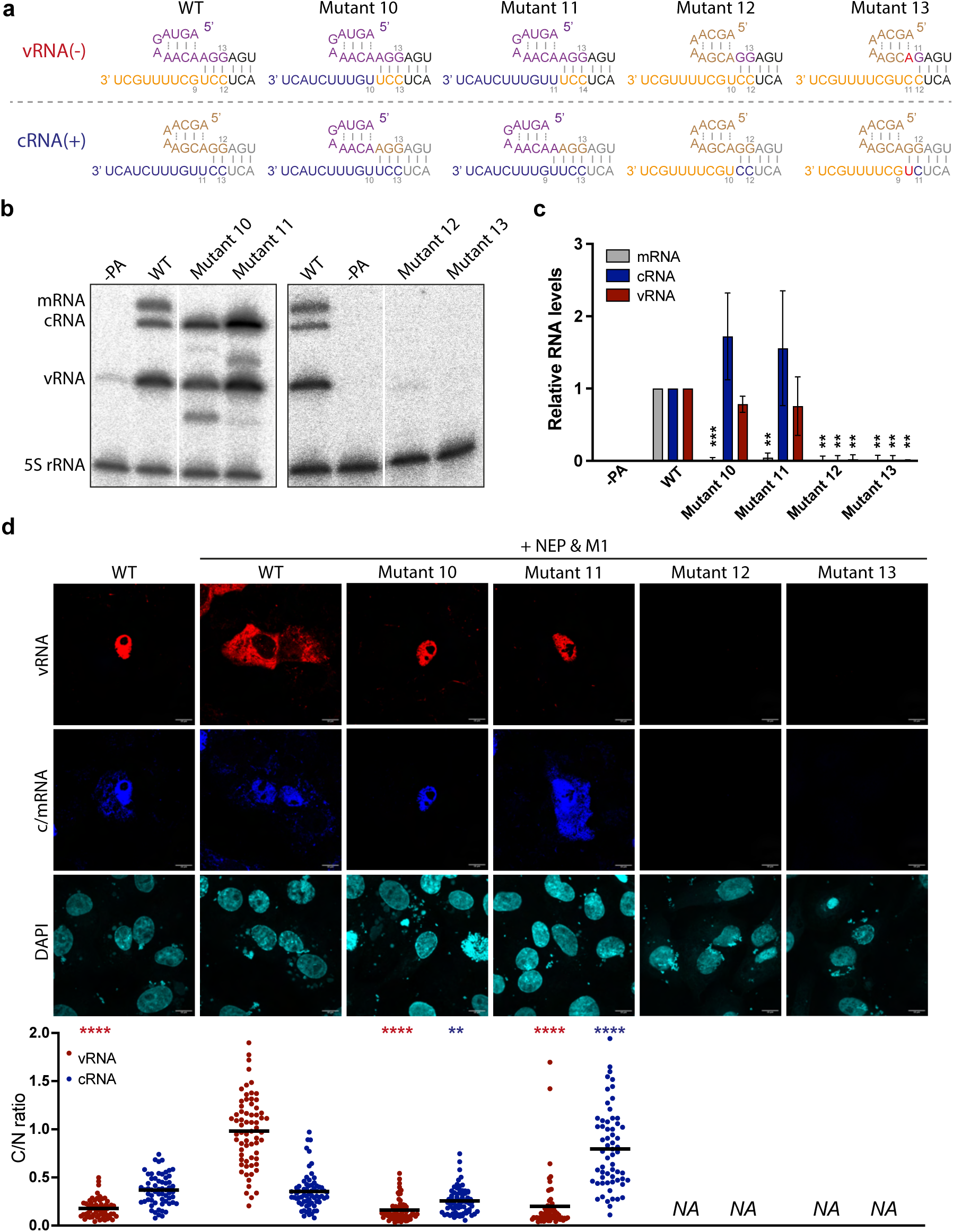
A nine-nucleotide cRNA-like 3′ promoter sequence enables cRNP export in a supportive promoter context. **a** Schematic representation of the wild-type and mutant vRNA and cRNA promoter sequences. Sequences are colour-coded as in Fig. 2a, and mutations are highlighted in red. **b** Primer extension analysis of viral RNAs from RNP reconstitution experiments using wild-type and mutant NA vRNA templates (WT and mutants 10-13). **c** Quantification of mRNA, cRNA, and vRNA levels from **b**. **d** Representative smFISH images and quantification of the cytoplasmic-to-nuclear (C/N) mean fluorescence intensity ratio from individual cells across two biological replicates: WT (n = 61), WT (+NEP&M1) (n = 67), mutant 10 (n = 70), and mutant 11 (n = 61). Experimental procedures and statistical analyses were performed as described in Fig. 2.

We first examined RNA synthesis by reconstituting RNPs with these mutant templates in the absence of NEP and M1. Mutants 10 and 11 remained replication competent but failed to produce detectable levels of mRNA (Fig. 5b, c). Comparable results were obtained in the presence of NEP and M1 (Supplementary Fig. 3). In contrast, mutants 12 and 13 did not support detectable RNA synthesis under either condition (Fig. 5b, c and Supplementary Fig. 3), indicating that a vRNA-like 5′ promoter is required for polymerase activity.

We next examined RNA localisation by smFISH. Mutant 10, which contains identical vRNA-like 5′ promoters and ten-nucleotide cRNA-like single-stranded 3′ promoter regions in both vRNA and cRNA, exhibited complete nuclear retention of both vRNA and cRNA (Fig. 5d). In contrast, mutant 11, in which the single-stranded regions of the vRNA and cRNA 3′ promoters are eleven-and nine-nucleotide long, respectively, showed efficient export of cRNA, whereas vRNA remained confined to the nucleus (Fig. 5d). Mutants 12 and 13 failed to produce sufficient levels of RNA for reliable quantification of the nuclear and cytoplasmic RNA signals.

Taken together, these findings demonstrate that a nine-nucleotide single-stranded 3′ promoter region is required for efficient RNP nuclear export. Extending the vRNA 3′ promoter to ten or eleven nucleotides abolished vRNP export, whereas shortening the cRNA 3′ promoter to nine nucleotides was sufficient to confer cRNP export competence. In contrast, a ten-nucleotide 3′ promoter failed to support export. Overall, these findings identify the length of the single-stranded 3′ promoter region as a major determinant of RNP nuclear export, while indicating that the 5′ promoter influences, but is not sufficient to specify, export competence.

### Exported cRNPs are packaged into virus-like particles

Our results above demonstrated that promoter architecture determines whether an RNP is retained in the nucleus or exported to the cytoplasm. We next asked whether nuclear export is sufficient to confer packaging competence or whether additional features unique to vRNPs are required for incorporation into virus particles.

To address this question, we established a virus-like particle (VLP) assay to examine packaging of export-competent cRNPs. Cells were transfected with expression plasmids encoding all viral proteins required for particle assembly together with a plasmid expressing a single vRNA segment (Fig. 6a). Only one vRNA segment was included in the assay to eliminate constraints imposed by intersegment packaging interactions and thereby assess the intrinsic packaging competence of individual RNP species. VLPs released into the culture supernatant were purified by haemadsorption to chicken red blood cells, a procedure that enriches intact particles while removing contaminating RNA released from damaged or lysed cells. RNA associated with purified VLPs was subsequently analysed by primer extension.

**Figure 6.**
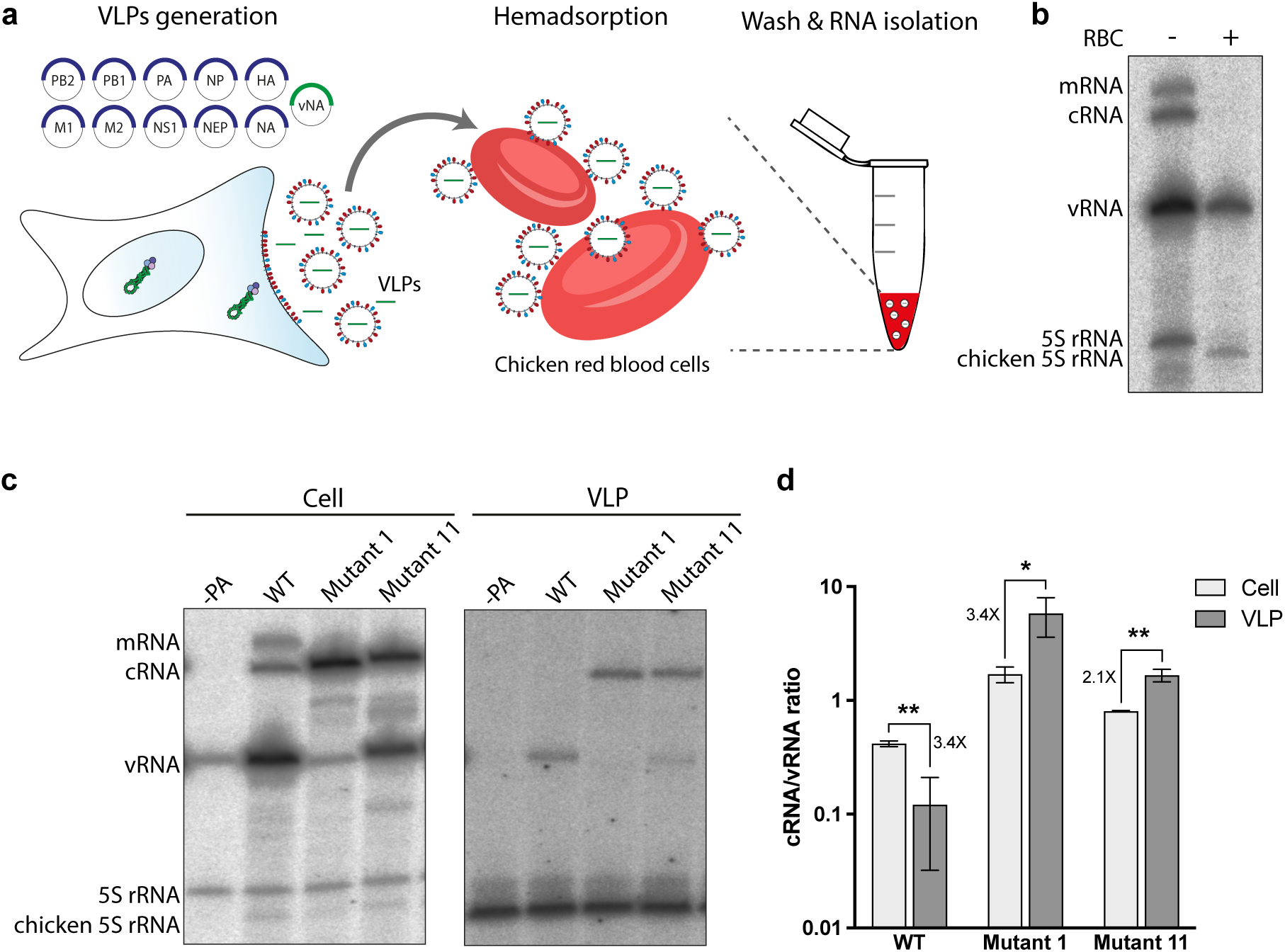
Export-competent cRNPs are incorporated into virus-like particles. **a** Schematic illustration of the virus-like particle (VLP) assay and haemadsorption procedure. **b** Validation of the haemadsorption procedure showing removal of contaminating RNA from a viral stock before and after binding to chicken red blood cells (RBCs). Viral RNA species were analysed by primer extension. **c** Primer extension analysis of viral RNAs from VLP experiments using wild-type and mutant NA vRNA templates. Total RNA from transfected cells (left panel) and RNA isolated from purified VLPs (right panel) were analysed. PA was omitted as a negative control. **d** Quantification of cRNA and vRNA levels from **c**. Relative RNA levels are presented as the cRNA/vRNA ratio, with fold changes indicated (mean ± SD, n = 3). Statistical significance was determined using Student’s t-test.

To validate the assay, we first analysed supernatants from influenza virus infected cells. Prior to purification, vRNA, cRNA and mRNA were readily detectable. Following haemadsorption, however, only vRNA remained detectable (Fig. 6b), confirming that the purification procedure selectively recovers RNA associated with intact virus particles and effectively removes non-particle-associated RNA.

We next analysed VLPs generated using wild-type and mutant promoters. As expected, VLPs produced with the wild-type promoter contained only vRNA, indicating selective incorporation of vRNPs (Fig. 6c, d). In contrast, mutants 1 and 11, both of which support efficient nuclear export of cRNPs, gave rise to VLPs containing readily detectable cRNA. Thus, cRNPs that acquire export competence through promoter modification are no longer excluded from particle incorporation.

These findings demonstrate that export competence and packaging competence are closely linked. Once cRNPs are rendered export competent, they are no longer excluded from particle incorporation. Together, these results suggest that selective genome packaging is initiated before virion assembly through differential nuclear export of RNP species and that subsequent cytoplasmic packaging mechanisms act to further refine genome selection.

### vRNA and cRNA promoter binding is associated with distinct polymerase conformational landscapes

Having identified promoter architecture as the determinant of export and packaging competence, we next sought to determine how the viral polymerase distinguishes between vRNA and cRNA promoters.

To address this question, we determined cryo-EM structures of influenza A virus polymerase in complex with NEP and either the 3′ vRNA or 3′ cRNA promoter. Purified polymerase (A/turkey/Turkey/1/2005) was incubated with NEP (A/WSN/33) and mix with either a 3′ vRNA (5′-GGCCUGCUUUUGCU-3′) or 3′ cRNA (5′-GGCCUUGUUUCUACU-3′) promoter before grid preparation and cryo-EM analysis.

Three-dimensional classification of both datasets identified encapsidase, replicase, and polymerase core-only conformations, with NEP bound at the PA-C/PB1-N interface as previously described^16^ (Supplementary Fig. 4 and 5). However, the distribution of promoter-bound states differed between the two datasets. The 3′ vRNA was observed in all three conformations, whereas the 3′ cRNA was detected only in the replicase and polymerase core-only conformations. Despite extensive three-dimensional classification, no cRNA-bound encapsidase population was identified. An apo, NEP-bound encapsidase population was present in both datasets, indicating that the polymerase-NEP complex itself is compatible with the encapsidase conformation.

The 3′ vRNA-bound encapsidase structure was refined to 2.71 Å resolution (Fig. 7a), with density for six promoter nucleotides resolved at the mode B surface site (Fig. 7b). U1 is sandwiched between the PB1 thumb domain and a PA-C β-sheet (residues 482–527), and the RNA extends towards the PB2 N-terminus, where it is stabilised by the PB2 627 domain. The 3′ cRNA-bound polymerase core-only structure was refined to 2.58 Å resolution (Fig. 7c) and similarly showed density for six promoter nucleotides at the mode B surface site (Fig. 7d). Structural alignment showed that the resolved cRNA nucleotides adopt essentially the same position across the polymerase surface as the vRNA nucleotides. Within this region, the two promoters differ in sequence only at positions 3 and 5, corresponding to guanine and uracil in vRNA and adenine and cytosine in cRNA, respectively.

**Figure 7.**
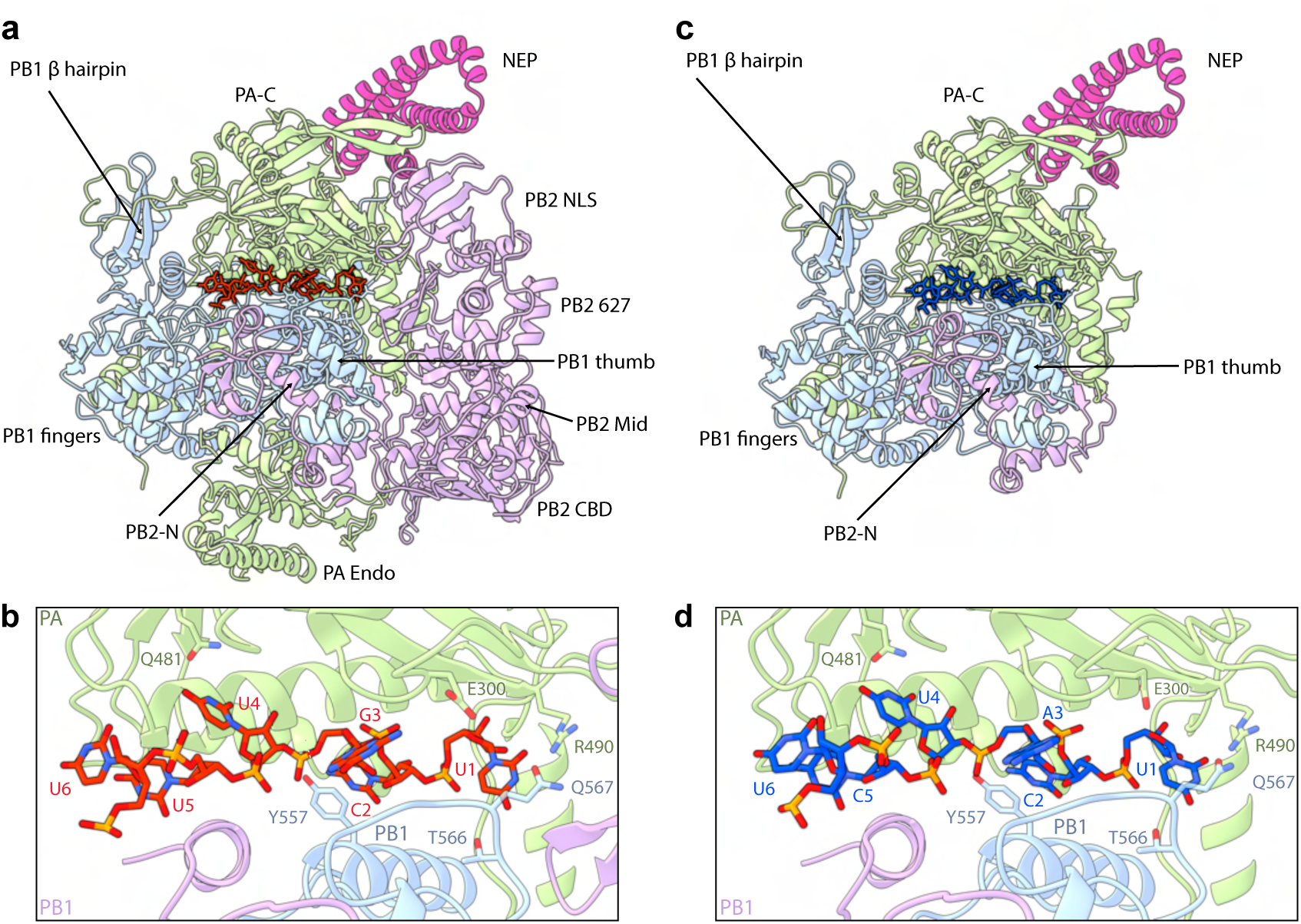
Cryo-EM analysis of NEP-bound influenza A virus polymerase complexes containing either a 3′ vRNA or 3′ cRNA promoter. **a** Cartoon representation of the viral polymerase in the encapsidase conformation with NEP bound to the PA-C/PB1 interface. **b** Six nucleotides of the 14-nucleotide 3′ vRNA promoter are resolved (red). **c** Cartoon representation of the viral polymerase core conformation with NEP bound at the PA-C/PB1 interface. **d** Six nucleotides of the 14-nucleotide 3′ cRNA promoter are resolved (blue).

Thus, despite occupying the same RNA-binding site on the polymerase surface, the 3′ vRNA and cRNA promoters are associated with distinct polymerase conformational landscapes (Supplementary Fig. 6). Both promoters were observed in the replicase and polymerase core-only states, whereas the RNA-bound encapsidase state was observed only with the 3′ vRNA promoter. Because apo, NEP-bound encapsidase particles were present in both datasets, the absence of a cRNA-bound encapsidase population suggests that the 3′ cRNA promoter does not stabilise, or may disfavour, this state. The sequence differences at positions 3 and 5 provide a potential basis for promoter discrimination, although whether they contribute to the distinct conformational landscapes remains unclear. These findings suggest that promoter identity can influence polymerase conformation, providing a potential mechanism linking promoter architecture to export competence.

## Discussion

Our study identifies selective nuclear export as a critical determinant of influenza virus genome packaging. Using direct RNA imaging, targeted promoter mutagenesis, functional VLP assays, and cryo-EM, we show that export competence is encoded within the viral RNA promoter. Specifically, we identify the promoter as the principal determinant of influenza virus RNP nuclear export. Exchange of the terminal promoter sequences completely reversed the export phenotype of vRNPs and cRNPs, demonstrating that export competence is encoded by promoter identity rather than RNA polarity. Subsequent dissection of the promoter showed that the critical determinants reside predominantly within the 3′ promoter, both the length and sequence of the single-stranded 3′ region contributing to export competence.

Importantly, neither the length nor the sequence of the 3′ promoter alone is sufficient to explain the export phenotypes. Although a nine-nucleotide single-stranded 3′ promoter was required for efficient export, shortening the cRNA promoter to nine nucleotides conferred only partial export in some promoter contexts. Likewise, substitution of the vRNA-specific 3′ promoter sequence reduced, but did not abolish, vRNP export, demonstrating that promoter sequence enhances rather than specifies export competence. Comparison of mutants 3, 5 and 11 further suggests that efficient export depends on the overall architecture of the promoter rather than any individual promoter feature. These mutants all possess an identical nine-nucleotide single-stranded cRNA-like 3′ promoter, either in the vRNA (mutant 5) or cRNA context (mutants 3 and 11), yet only mutant 11 supported efficient export. The principal structural distinction is that mutant 11 forms an additional base pair within the promoter duplex, raising the possibility that promoter duplex formation contributes to stabilising an export-competent promoter conformation. Together, these findings suggest that nuclear export depends on recognition of a specific promoter architecture, in which the length and sequence of the single-stranded 3′ promoter, together with the sequence of the 5′ promoter and the extent of promoter duplex formation, cooperate to generate an export-competent RNP.

These findings provide a mechanistic explanation for the long-standing observation that cRNPs are excluded from virions despite their close structural and compositional similarity to vRNPs^5–7^. Our data show that cRNPs are selectively retained in the nucleus, indicating that discrimination between vRNPs and cRNPs occurs prior to virion assembly. Current models of influenza genome packaging largely focus on events occurring in the cytoplasm, including segment-specific RNA-RNA interactions and assembly of the characteristic 7+1 genome bundle^8,9^. While these mechanisms undoubtedly contribute to genome selection, our findings indicate that only a subset of viral RNPs gains access to the cytoplasmic assembly pathway in the first place. Thus, rather than relying exclusively on cytoplasmic selection mechanisms, influenza virus employs nuclear export as an early checkpoint to enforce genome specificity.

The VLP experiments further support this view. In particular, promoter mutants that confer cRNP export competence also permit cRNA incorporation into virus-like particles. In contrast, wild-type cRNPs remain nuclear-retained and are excluded from packaging. These observations demonstrate a direct link between export competence and packaging competence and suggest that selective genome packaging is initiated through differential nuclear export of RNP species. Cytoplasmic packaging mechanisms may subsequently refine this selection process, but the decision of which RNPs are eligible for packaging appears to be made earlier, within the nucleus.

The promoter mutagenesis experiments further illustrate how promoter architecture coordinates multiple aspects of viral RNA biology, including polymerase activity. We show that polymerase activity requires at least one promoter to retain a vRNA-like 5′ promoter sequence, as templates in which both promoters contain cRNA-like 5′ sequences are inactive in transcription and replication (mutants 7, 12, and 13). Interestingly, many promoter mutants remained replication competent despite exhibiting severe defects in transcription, indicating that the requirements for replication and transcription are not equivalent. In particular, extension of the single-stranded 3′ promoter region strongly impaired mRNA synthesis while allowing substantial accumulation of cRNA and vRNA. These observations suggest that cap-dependent transcription initiation imposes more stringent structural requirements on promoter architecture than de novo replication.

Several promoter mutants exhibited increased accumulation of cRNA (mutants 1, 5, 8, and 9). One possibility is that promoter configurations that fail to support efficient transcription favour polymerase states associated with replication. Although the present study was not designed to investigate the mechanistic basis of replication-transcription switching, the increased accumulation of cRNA observed for several promoter mutants suggests that promoter architecture contributes to regulating RNA synthetic outcome in addition to export competence. Thus, the influenza virus promoter functions as a multifunctional regulatory element that coordinates transcription, replication and intracellular trafficking.

NEP has previously been shown to regulate polymerase activity in a dose-dependent manner^16,31^. Consistent with these observations, inclusion of NEP and M1 in our smFISH assays resulted in a modest reduction in transcription, whereas replication remained largely unaffected. Importantly, the relative levels of vRNA and cRNA detected by smFISH closely mirrored those measured by primer extension analysis.

Our findings also provide a mechanistic framework for interpreting earlier observations that promoter mutations can promote cRNA packaging^6^. In that study, cRNA incorporation into virions was analysed in the context of a promoter-up background, making it difficult to distinguish effects on promoter activity from effects on packaging specificity. Although those experiments demonstrated that promoter architecture can influence packaging, the underlying basis for this phenotype remained unresolved. By combining direct RNA imaging, promoter mutagenesis and VLP assays, we show that promoter architecture regulates export competence and that acquisition of export competence is sufficient to permit RNP incorporation.

The cryo-EM analyses provide a possible structural basis for understanding how promoter architecture is sensed by the viral polymerase. Both the 3′ vRNA and cRNA promoters bind the same mode B surface site and adopt similar local binding modes. Nevertheless, the two promoters are associated with distinct polymerase conformational landscapes. Whereas the 3′ vRNA dataset contained encapsidase, replicase and core-only polymerase conformations bound to 3′ vRNA, the corresponding 3′ cRNA dataset contained only replicase and core-only polymerase conformations bound to 3′ cRNA and, importantly, lacked the RNA-bound encapsidase conformation. These observations demonstrate that promoter identity influences polymerase conformation despite engagement of a common RNA-binding site. However, the structural analyses do not establish a direct causal relationship between a specific polymerase conformation and nuclear export. Rather, they show that promoter architectures associated with distinct biological outcomes are also associated with distinct polymerase structural states. Thus, promoter-dependent modulation of polymerase conformation provides a plausible mechanism through which export competence may ultimately be regulated.

Several limitations should be considered. Our analyses were performed primarily in reconstituted RNP and virus-like particle systems rather than during complete virus infection. In addition, the cryo-EM structures capture isolated promoter-bound polymerase complexes and do not include the complete export machinery. Consequently, the molecular events linking promoter recognition to CRM1/XPO1-dependent export remain unresolved. Future studies aimed at determining structures of complete export complexes containing promoter RNA, polymerase, NEP and host export factors will be required to define the underlying molecular mechanism.

Together, our results demonstrate that the two promoter termini act in a coordinated manner to control viral RNA fate. Disruption of the nine-nucleotide single-stranded 3′ region, alteration of key promoter residues, or modification of the 5′ promoter sequence impairs export competence without necessarily abolishing replication, highlighting promoter architecture as a multifunctional element that links RNA synthesis to RNP trafficking. We propose that influenza virus exploits its nuclear replication strategy to impose a promoter-encoded checkpoint that restricts non-genomic RNPs from accessing the cytoplasmic assembly pathway. In this model, only RNPs possessing an export-competent promoter architecture are licensed for nuclear export and subsequent packaging. Our findings reveal that genome selection begins before virion assembly and establish nuclear export as an integral component of the influenza A virus genome-packaging pathway.

---

**Supplementary Figure 1.**
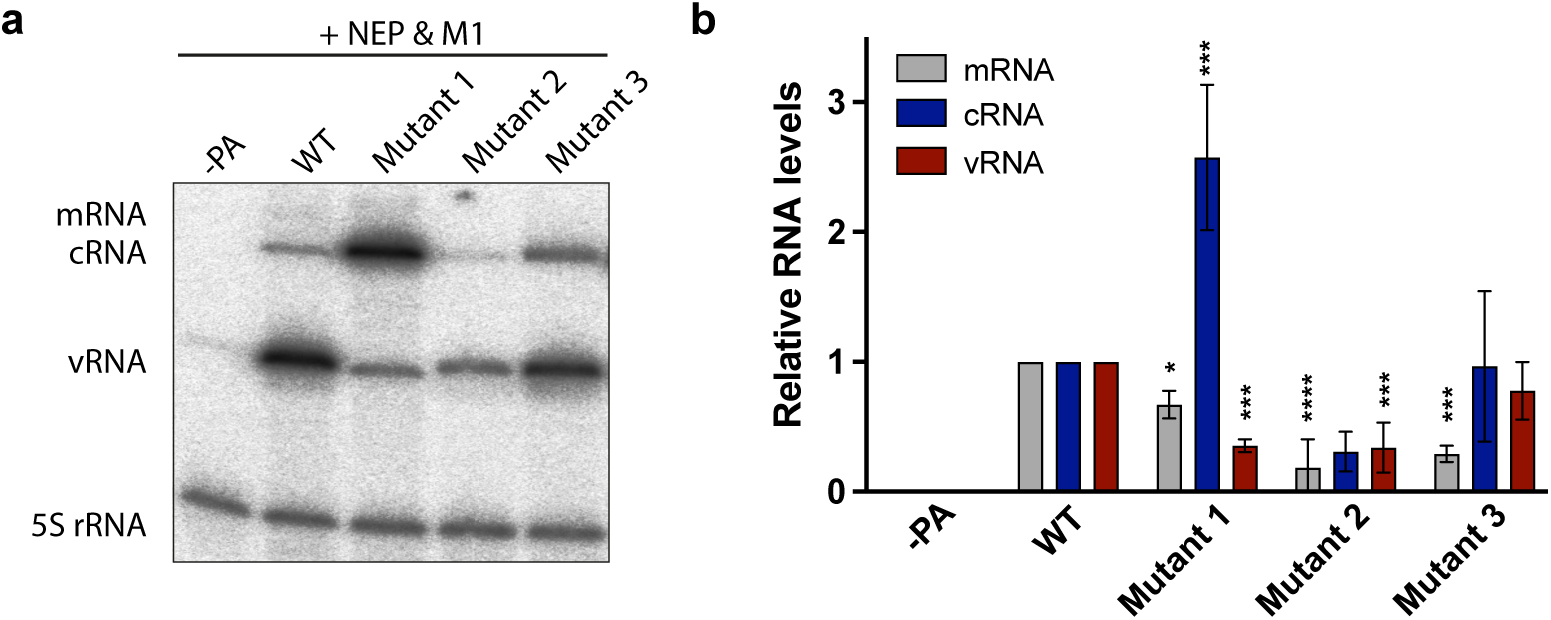
**a** Primer extension analysis of viral RNAs from RNP reconstitution experiments in the presence of NEP and M1 using wild-type and mutant NA vRNA templates (WT and mutants 1-3). PA was omitted as a negative control. **b** Quantification of mRNA, cRNA, and vRNA levels from (A). RNA levels were normalised to the WT condition (mean ± SD, n = 3). Statistical analysis was performed using one-way ANOVA with multiple comparisons relative to WT.

**Supplementary Figure 2.**
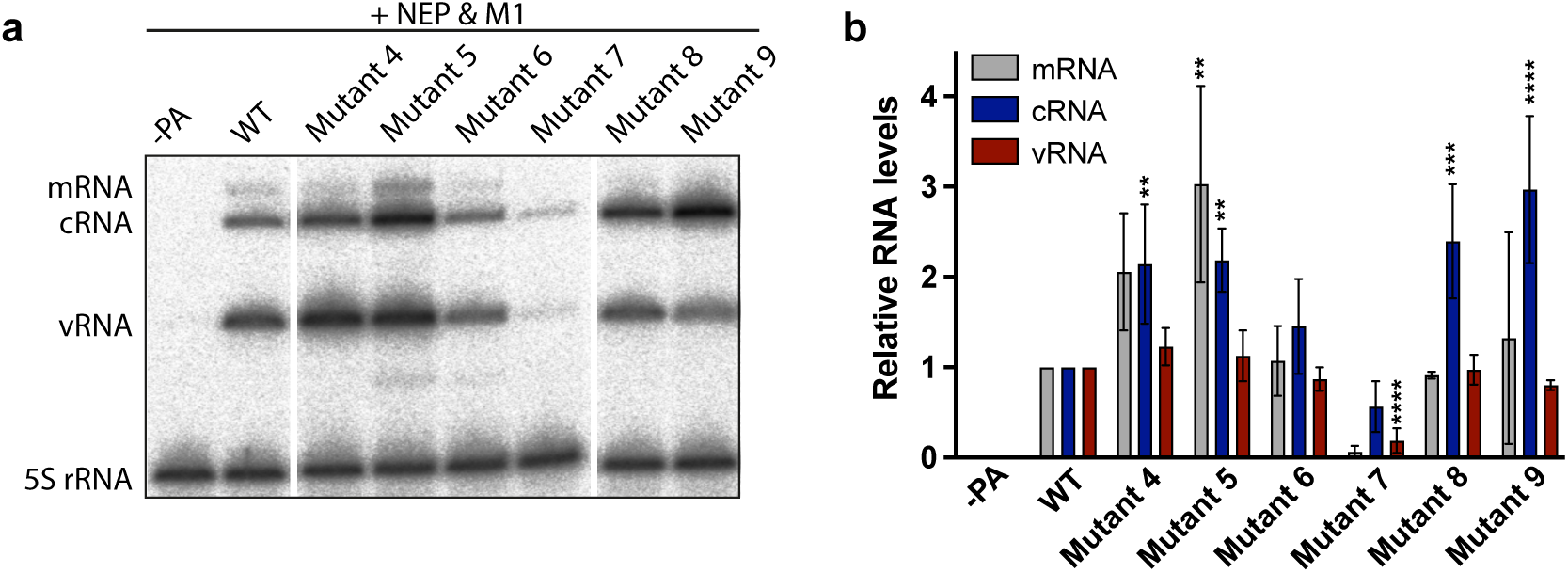
**a** Primer extension analysis of viral RNAs from RNP reconstitution experiments in the presence of NEP and M1 using wild-type and mutant NA vRNA templates (WT and mutants 4-9). PA was omitted as a negative control. **b** Quantification of mRNA, cRNA, and vRNA levels from **a**. RNA levels were normalised to the WT condition (mean ± SD, n = 3). Statistical analysis was performed using one-way ANOVA with multiple comparisons relative to WT.

**Supplementary Figure 3.**
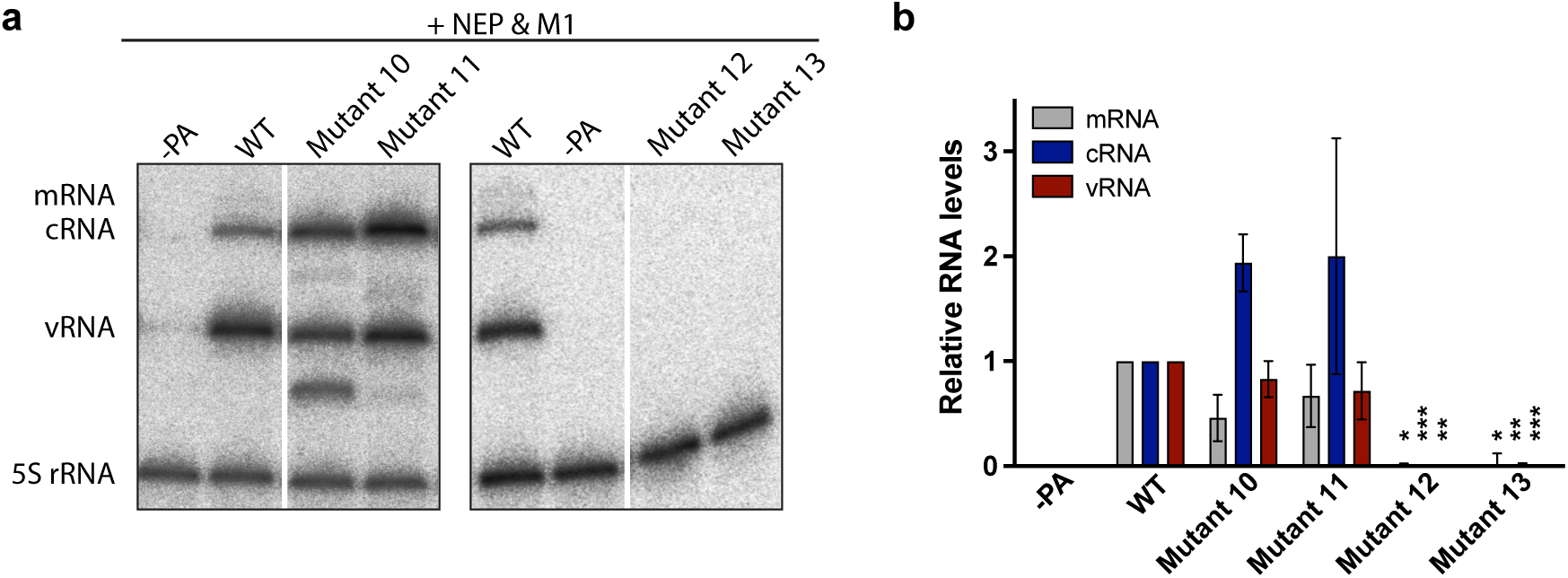
**a** Primer extension analysis of viral RNAs from RNP reconstitution experiments in the presence of NEP and M1 using wild-type and mutant NA vRNA templates (WT and mutants 10-13). PA was omitted as a negative control. **b** Quantification of mRNA, cRNA, and vRNA levels from **a**. RNA levels were normalised to the WT condition (mean ± SD, n = 3). Statistical analysis was performed using one-way ANOVA with multiple comparisons relative to WT.

**Supplementary Figure 4.**
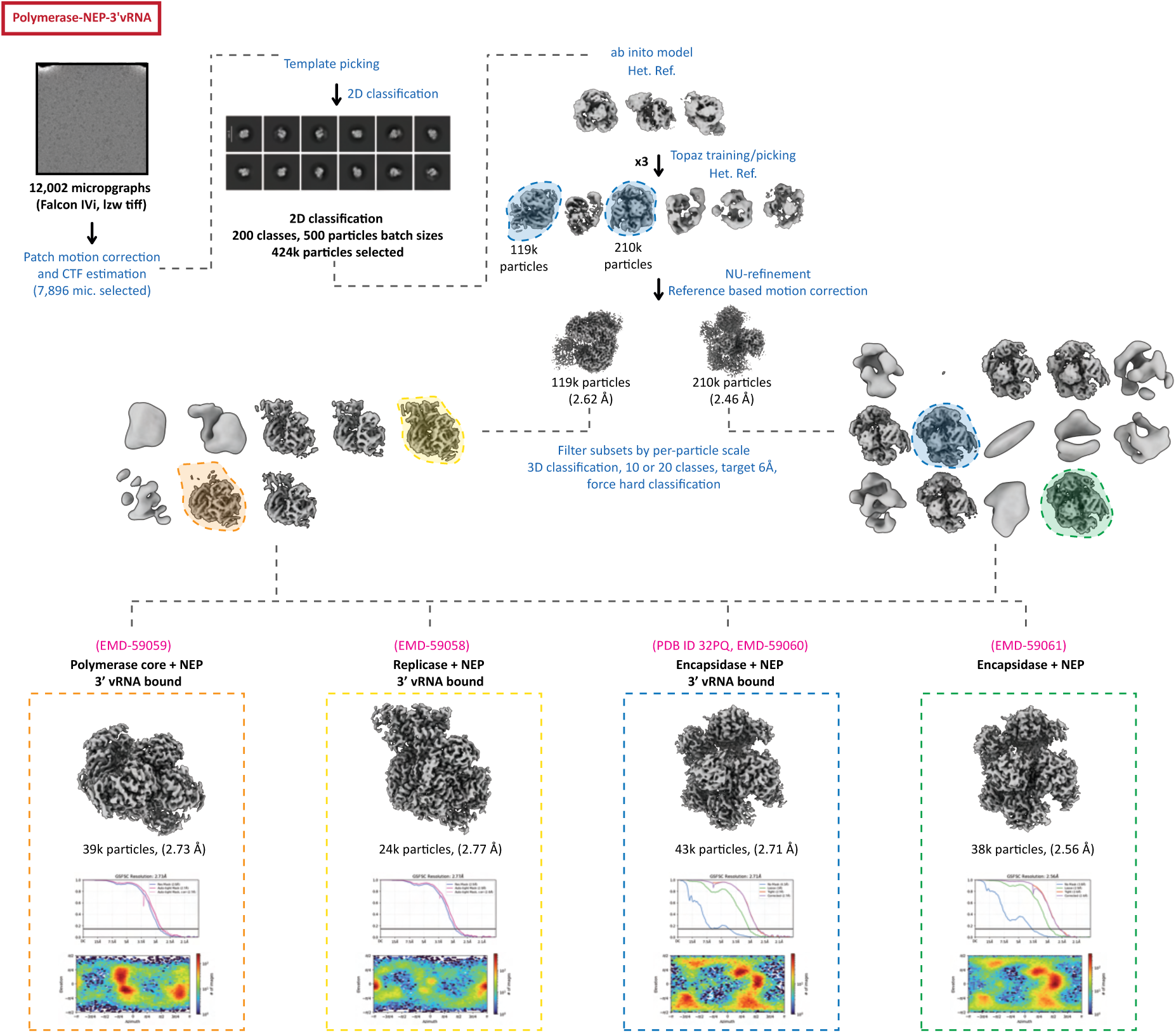
Cryo-EM data collection and processing workflow for influenza A virus polymerase-NEP complexes in the presence of the 3’ vRNA promoter.

**Supplementary Figure 5.**
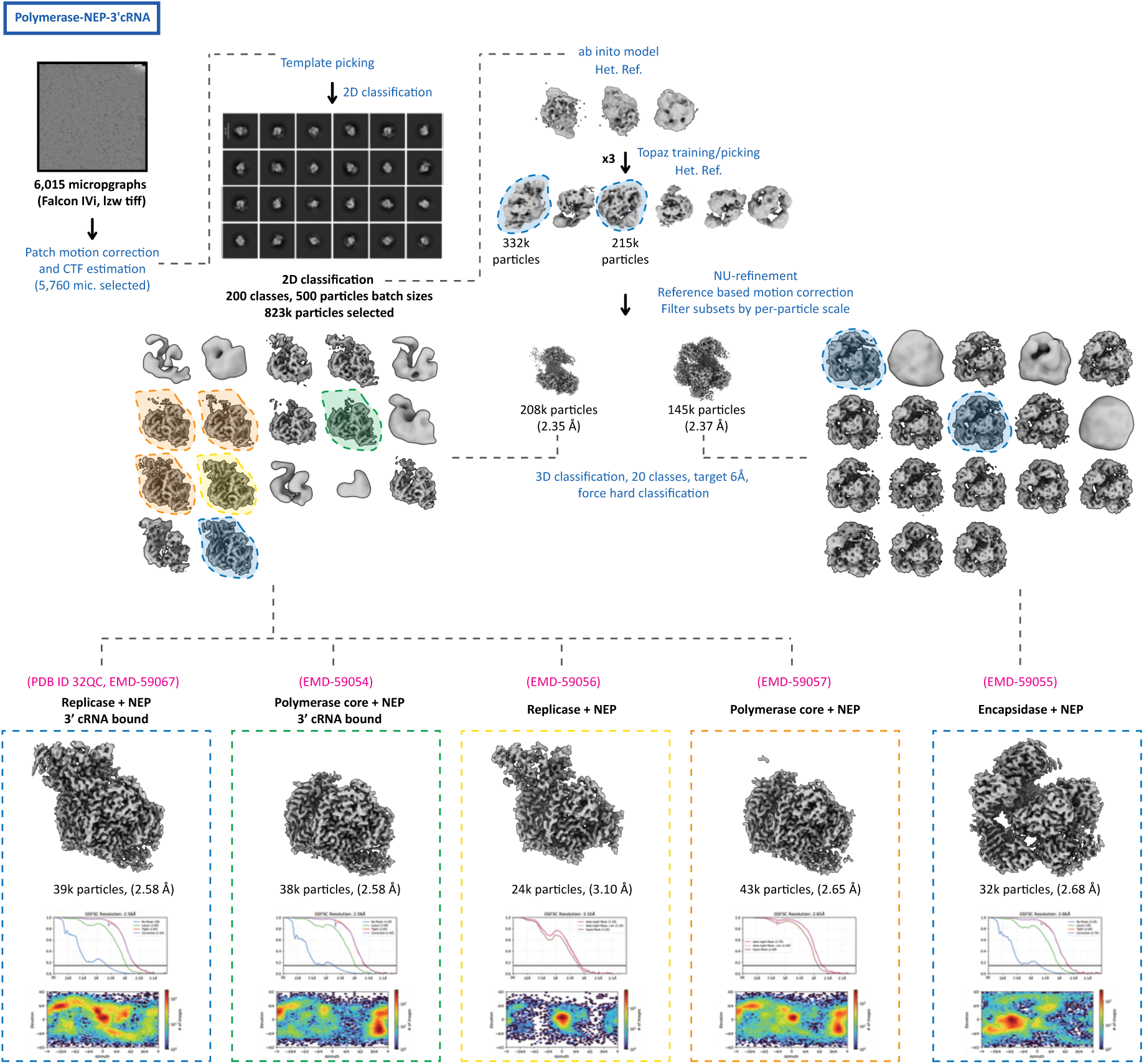
Cryo-EM data collection and processing workflow for influenza A virus polymerase-NEP complexes in the presence of the 3’ cRNA promoter.

**Supplementary Figure 6.**
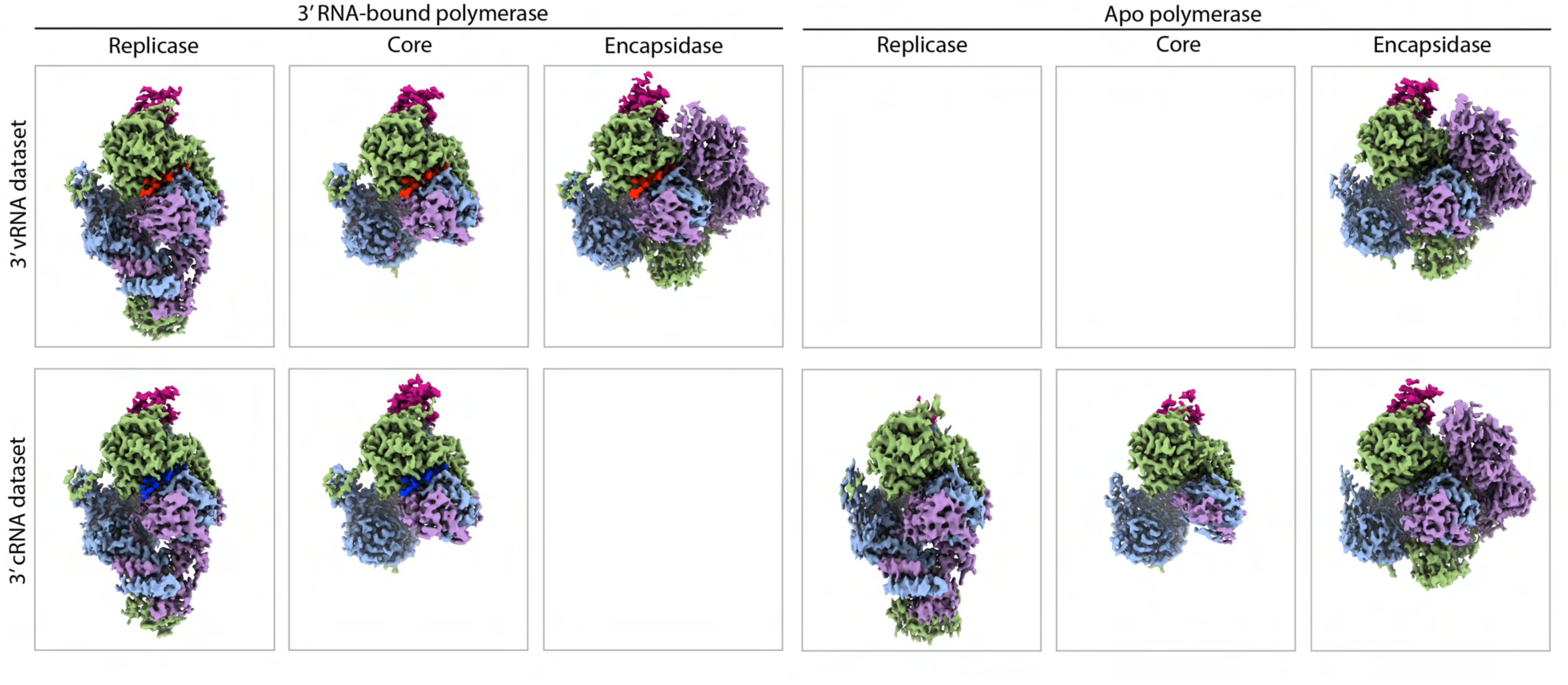
Cryo-EM density maps of influenza A virus polymerase-NEP complexes showing replicase, polymerase core-only, and encapsidase conformations in the 3′ vRNA dataset (upper panel) and 3′ cRNA dataset (lower panel), either bound to 3′ promoter RNA (left) or without RNA (apo; right).

## Materials and methods

### Cells

HEK-293T cells and Vero E6 cells were cultured in Dulbecco′s Modified Eagle′s Medium (DMEM; Sigma) supplemented with 10% foetal bovine serum (FBS; Sigma) and maintained at 37°C in a humidified atmosphere containing 5% CO_2_. All cell lines were obtained from the Cell Bank of the Sir William Dunn School of Pathology, University of Oxford.

### Plasmids

Plasmids pCAGGS-PB2, pCAGGS-PB1, pCAGGS-PA, and pCAGGS-NP, encoding proteins from influenza A/WSN/33 (H1N1), were kindly provided by Adolfo García-Sastre. Plasmids pCAGGS-NEP, pCAGGS-M1, pcDNA-NS1, and pPOLI-NA-RT, also encoding sequences from A/WSN/33, were generated as previously described^16,32,33^. Site-directed mutagenesis was used to introduce mutations into the promoter region of vRNA encoded by the pPOLI-NA-RT plasmid. The endonuclease-inactive mutant plasmid pCAGGS-PA-D108A was generated by PCR amplification of the PA coding sequence from pcDNA-PA-D108A^30^, followed by Gibson assembly into the pCAGGS vector. The pcDNA-M2 expression plasmid was generated by PCR amplification of the M2 coding sequence from pPOLI-M-RT^32^, followed by cloning into pcDNA using BamHI and XhoI restriction sites. Plasmids pCAGGS-PR8-HA and pCAGGS-PR8-NA, encoding HA and NA from influenza A/PR8/34 (H1N1), were generated by PCR amplification of the corresponding genes from pPOLI-PR8-HA and pPOLI-PR8-NA^34^, followed by cloning into pCAGGS using NotI and XhoI restriction sites.

### RNP reconstitution assay

HEK-293T cells seeded in 12-well plates were transiently transfected with 100 ng each of pCAGGS-PB2, pCAGGS-PB1, and pCAGGS-PA (wild-type or the endonuclease-inactive mutant PA D108A), together with 200 ng of pCAGGS-NP and 100 ng of pPOLI-NA-RT (wild-type or mutant constructs). Transfections were performed using Lipofectamine 2000 (Invitrogen) according to the manufacturer′s instructions. At 24 h post-transfection, cells were harvested either for total RNA extraction and primer extension analysis or for immunoblotting.

### RNA isolation and primer extension analysis

Total RNA was extracted using TRI reagent (Sigma) according to the manufacturer′s instructions and analysed by primer extension as previously described^35^. Briefly, 2 μg of total RNA was reverse transcribed using Superscript III reverse transcriptase and ^32^P-labelled, strand-specific primers targeting the NA segment. A primer specific for 5S rRNA was included as a loading control^30^. cDNA products were resolved on 6% denaturing polyacrylamide gels containing 7 M urea. Radioactive signals were detected using phosphor screens and visualised with an FLA-5000 scanner (Fuji). Images were analysed using Fuji-ImageJ software.

### Immunoblotting

Total cell lysates were prepared in RIPA buffer (50 mM Tris-HCl, pH 7.4, 150 mM NaCl, 1% NP-40, 0.5% sodium deoxycholate, and 0.1% SDS) supplemented with 1× cOmplete protease inhibitor cocktail (Roche) and analysed by western blotting. Primary antibodies against PA (GTX125932, GeneTex), NP (GTX125952, GeneTex), and GAPDH (2118, Cell Signaling Technology) were used, followed by IRDye-conjugated goat anti-rabbit or goat anti-mouse secondary antibodies (LI-COR).

### smFISH assay of viral RNA localisation

All probes were designed and labelled as previously described^16^. Probes targeting the NA segment mRNA/cRNA were designed using the Stellaris Probe Designer (Table S1)^36^. As for the vRNA probes, a 28-nucleotide overhang sequence (5′ACACTCGGACCTCGTCGACATGCATTAA-3′) was added to the 5′ end of each probe. Fluorescently labelled detection oligonucleotides complementary to the overhang sequence were used for probe visualisation: an ATTO550-labelled oligonucleotide used for vRNA detection, and an ATTO650-labelled oligonucleotide was used for mRNA/cRNA detection. Vero E6 cells were seeded onto 13 mm glass coverslips in 24-well plates and transfected with 100 ng each of pCAGGS-PB2, pCAGGS-PB1, and pCAGGS-PA (wild-type or the endonuclease-inactive mutant PA D108A), 200ng of pCAGGS-NP, 30ng of pPOLI-NA-RT (wild-type or mutants), 100ng of pCAGGS-GFP-NEP, and 50ng of pCAGGS-M1 using Lipofectamine 3000 following the manufacturer′s instructions. At 24 h post-transfection, cells were fixed with 4% paraformaldehyde for 20 min at room temperature and permeabilised in 70% ethanol overnight at 4 °C. The following day, cells were washed with PBS to remove residual ethanol, incubated in 80% formamide at 37 °C for 10 min, and rehydrated in 2× SSC buffer (30 mM sodium citrate, pH 7.0, 300 mM NaCl) at room temperature for 10 min. Hybridisation was performed at 37 °C for 4 h using freshly prepared hybridisation buffer containing 100 nM vRNA probes, 300 nM m/cRNA probes, 2× SSC, 10% formamide, 10% dextran sulfate, and 2 mM ribonucleoside vanadyl complex. Following hybridization, cells were washed twice for 10 min in pre-warmed 2× SSC containing 10% formamide at 37 °C. Coverslips were mounted in mounting medium containing 4′,6-diamidino-2-phenylindole (DAPI) (Ibidi) for nuclear staining.

### Confocal microscopy and image analysis

Fluorescence images were collected on an FV3000 confocal laser-scanning microscope (Olympus) using a 40× objective and HSSD/SD detectors. Excitation was provided by 405 nm, 488 nm, 561 nm, and 630 nm laser lines to visualize DAPI, GFP, vRNA, and m/cRNA FISH signals, respectively. Images were acquired at a spatial resolution of 310 nm per pixel with a frame size of 1024 × 1024 pixels, covering a total z-volume of 6 µm sampled at 1.2 µm intervals.

Image processing and analysis were performed using ImageJ and Cellpose software. Nuclear boundaries were defined based on DAPI staining, while cellular boundaries were segmented using GFP fluorescence. FISH signals were quantified by measuring the mean fluorescence intensity within the defined nuclear and cytoplasmic regions, as previously described^16^.

### Virus-like particle assay for RNP packaging analysis

Virus-like particles (VLPs) were generated by transfecting HEK-293T cells with 500 ng each of pCAGGS plasmids encoding PB2, PB1, PA, HA, NP, NA, and NEP, together with 1,000 ng of pCAGGS-M1, 500 ng of pcDNA-NS1, 100 ng of pcDNA-M2, and 1,000 ng of pPOLI-NA-RT (wild-type or mutant constructs) using Lipofectamine 2000 according to the manufacturer′s instructions. At 6 h post-transfection, the medium was replaced with DMEM supplemented with 2% fetal bovine serum (FBS). Supernatants containing VLPs were harvested 48 h later, clarified by centrifugation at 10,000 × g for 5 min, and incubated with 1% chicken red blood cells (RBCs) in PBS for haemadsorption at 4 °C. After 1 h, RBCs with adsorbed VLPs were washed twice with PBS and centrifuged at 800 × g for 5 min at 4 °C to remove unbound material. The resulting RBC pellets were processed for total RNA isolation and primer extension analysis. In parallel, transfected HEK-293T cells were harvested for RNA isolation and primer extension analysis to serve as input controls.

### Protein expression and purification

The influenza A/turkey/Turkey/1/2005 (H5N1) polymerase complex was produced by co-expressing codon-optimized PB1, PB2, and PA subunits in Sf9 cells using the MultiBac system and purified as described previously^37^. The polymerase carries PA T97I and PA E349K substitutions, which are located at the interface of the polymerase symmetric dimer, preventing its formation and biasing the polymerase heterotrimer towards an encapsidase conformation^16,24,37^. It also carries the mammalian-adaptive substitutions PB2 E627K and PA N383D, together with PB2 D701N, PB2 K702R, and PB2 S714R, as well as PB1 K577E and PA Q556R, which were evolved from serial passaging in human cells lacking ANP32A and ANP32B^37,38^. Purification was performed by affinity chromatography using IgG Sepharose resin (GE Healthcare), followed by size-exclusion chromatography on a Superdex 200 Increase 10/300 GL column (GE Healthcare) equilibrated in SEC buffer (25 mM HEPES-NaOH, pH 7.5, 150 mM NaCl, 5% (v/v) glycerol, and 1 mM dithiothreitol).

NEP of A/WSN/33 was expressed from a pGEX-6-P-1 bacterial expression vector with an N-terminal GST tag, followed by a PreScission 3C protease cleavage site and a seven amino acid linker. Expression was performed in BL21 E. coli cells at 37 °C with shaking at 180 rpm. When cultures reached an OD_600_ of 0.6-0.8, protein expression was induced with 0.5 mM IPTG, and cells were incubated overnight at 18 °C with shaking at 180 rpm. Cell pellets were lysed in wash buffer containing 25 mM Tris-HCl (pH 7.5), 300 mM NaCl, 10% glycerol, 1 mM dithiothreitol, supplemented with 1 mg/ml lysozyme from chicken egg white (Sigma), 100 *μ*g/ml RNase A, and one protease inhibitor cocktail tablet (Roche) per 100 ml of resuspended culture. Cells were sonicated for 30 s followed by a 30 s pause; this cycle was repeated three times. The lysate was clarified by centrifugation at 35,000 × g for 1 h prior to incubation with Glutathione Sepharose 4B resin (GE Life Sciences) pre-equilibrated in wash buffer. The lysate was incubated with the resin for 1 h at 4 °C, after which the resin was washed three times with wash before cleavage with human rhinovirus (HRV) 3C protease (produced in-house) for 1 h at 4 °C. The resin was separated from the supernatant by centrifugation at 1500 × g for 5 min at 4 °C. The protein-containing supernatant was concentrated using a centrifugal concentrator (Millipore) with a 3 kDa molecular weight cut-off. Subsequent size-exclusion chromatography was performed on a Superdex 200 Increase 10/300 column (Cytiva) equilibrated in SEC buffer comprising 25 mM Tris-HCl (pH 7.5), 300 mM NaCl, 5% glycerol and 1 mM dithiothreitol.

### Cryo-EM grid preparation

To prepare cryo-EM samples, NEP was mixed with polymerase at a 5:1 molar ratio to a final concentration of 0.35 mg/ml in SEC buffer containing 230 mM NaCl. Prior to plunging the same sample was mix with 1.2 molar excess of either 3′ vRNA: 5′-GGCCUGCUUUUGCU-3′ or 3′ cRNA: 5′-GGCCUUGUUUCUACU-3′ promoter. UltraAufoil R1.2/1.3 300 mesh grids were glow-discharged on both side prior to application of 1.7 μl of samples on each side. Grids were prepared under 100% relative humidity and blotted for 3 s with a blot force of 0, followed by vitrification in liquid ethane using a Vitrobot Mark IV (Thermo Fisher Scientific).

### Cryo-EM image collection

Cryo-EM data were collected at the Oxford Particle Imaging Centre (OPIC) using a 300 kV G3i Titan Krios microscope (Thermo Fisher Scientific) equipped with a SelectrisX energy filter and Falcon IVi direct electron detector. Automated data collection was performed using EPU 3.10, and movies were recorded in lzw compressed tiff format. Data were collected using AFIS with a total dose of approximately 50 e^-^/Å^2^, a calibrated pixel size of 0.932 Å/pixel, and a 10 eV slit. Specific data collection parameters for the NEP-polymerase complexes containing either the 3′ vRNA or 3′ cRNA promoter are provided in Table S2.

### Cryo-EM data processing

All datasets were processed using CryoSPARC v4.5-v5.0.5. EER movies were fractionated into 60 frames without upsampling. Pre-processing was performed using Patch Motion Correction and Patch CTF Estimation with default settings. Motion-corrected micrographs with poor image quality or suboptimal CTF statistics were removed by manual curation.

For complexes containing either the 3′ vRNA or 3′ cRNA promoter, data were processed using the same workflow. Following motion correction and CTF estimation, particles were template-picked using previously determined encapsidase, replicase, and polymerase core-only 2D classes derived from PDB 8R1J processing. Following 2D classification, well-resolved classes were selected and used to generate three ab initio models further refined by heterogeneous refinement.

Particles belonging to high-resolution classes were used to train a Topaz model, which was subsequently used for particle picking across the full dataset. The resulting particles were subjected directly to heterogeneous refinement using the six previously generated ab initio models, each supplied twice as a reference. This sequence has been used iteratively to increase the number of particles. Separate NU refinements were performed for the encapsidase and replicase, and polymerase core-only classes, followed by reference-based motion correction and a second round of independent NU refinement.

To distinguish apo and RNA-bound NEP-polymerase complexes, 3D classification without alignment was performed on each class followed by a local NU refinement for each of the selected classes. Details are in included in Supplementary Fig 4 and 5.

### Structure determination and model refinement

Initial model building for the 3′ vRNA or 3′ cRNA promoter-bound NEP-polymerase complexes was performed using PDB entry 8R1J as a starting model. Initial fitting was carried out in UCSF ChimeraX using either the complete replicase or encapsidase conformation. Flexible fitting was subsequently performed in WinCoot 0.9.8.7^39^ using the restraints module with restraints generated at 4.3 Å resolution and chain refinement.

Multiple rounds of manual model adjustment in WinCoot followed by real-space refinement in PHENIX 2.0 were used to optimise model geometry and fit to the density. Final model quality and map-to-model agreement were assessed using MolProbity within PHENIX^40^. Statistics are summarized in Data Table S2. Structural analysis and figure preparation were performed using UCSF ChimeraX.

---

**Data Table S1:**
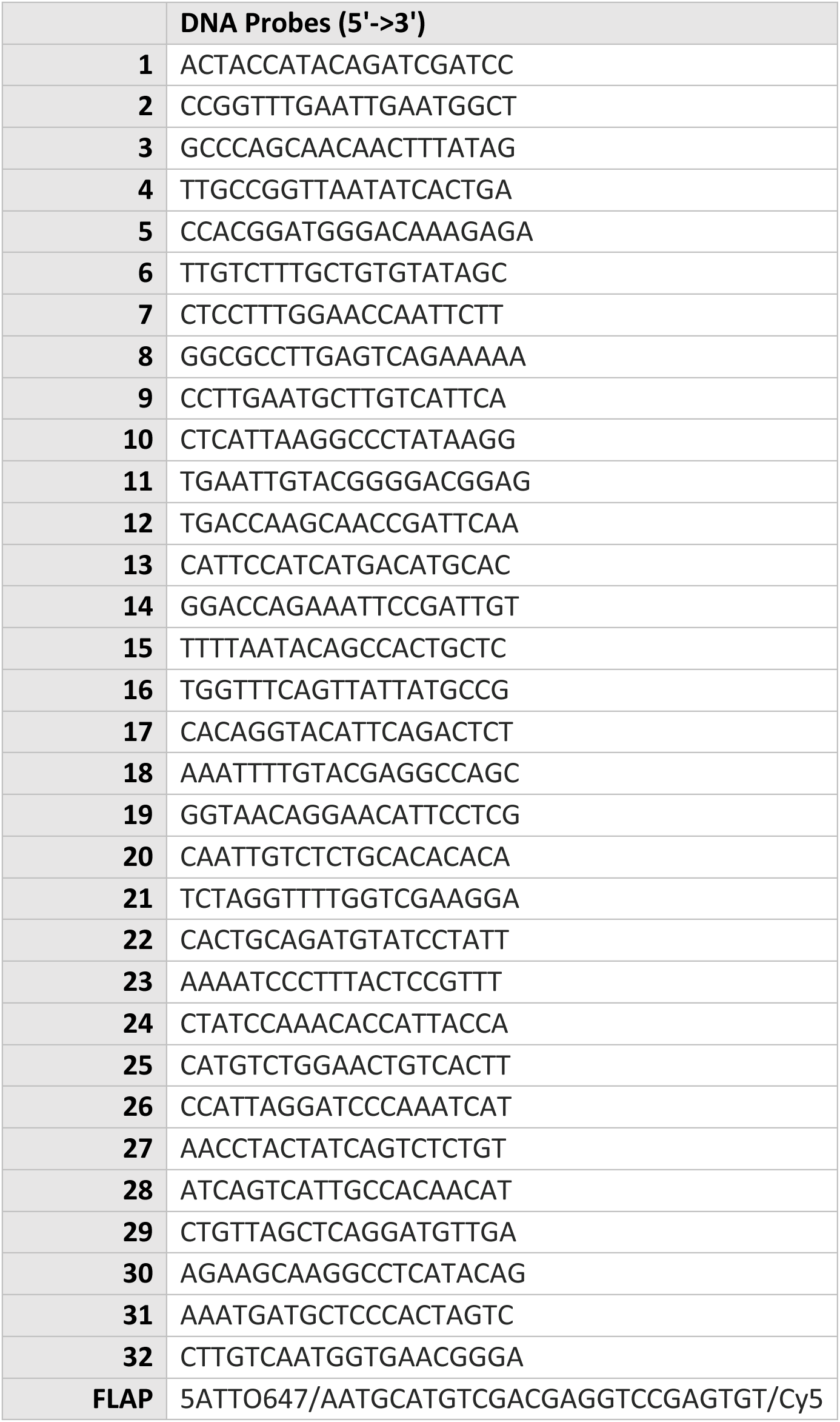
Probes against NA positive-sense RNA.

**Data Table S2:** Cryo-EM data collection and refinement parameters.

|  | Polymerase-NEP-3'cRNA dataset |  |  |  |  | PolymeraseI-NEP-3'vRNA dataset |  |  |  |
| --- | --- | --- | --- | --- | --- | --- | --- | --- | --- |
|  | 3' cRNA bound |  | NO RNA bound |  |  | 3' vRNA bound |  |  | NO RNA bound |
|  | Replicase | Core | Encapsidase | Replicase | Core | Replicase | Core | Encapsidase | Encapsidase |
|  | EMD-59067 | EMD-59054 | EMD-59055 | EMD-59056 | EMD-59057 | EMD-59058 | EMD-59059 | EMD-59060 | EMD-59061 |
|  | 32QC |  |  |  |  |  |  | 32PQ |  |
| **Data collection** |  |  |  |  |  |  |  |  |  |
| Microscope | Titan Krios OPIC |  |  |  |  | Titan Krios OPIC |  |  |  |
| Voltage (kV) | 300 |  |  |  |  | 300 |  |  |  |
| Detector | Falcon IVi - SelectrisX |  |  |  |  | Falcon IVi - SelectrisX |  |  |  |
| Recording mode | lzw tiff |  |  |  |  | lzw tiff |  |  |  |
| Magnification | 130,000 |  |  |  |  | 130,000 |  |  |  |
| Movie/micrograph pixel size (Å) | 0.932 |  |  |  |  | 0.932 |  |  |  |
| Dose rate (e-/px/sec) | 9.51 |  |  |  |  | 9.51 |  |  |  |
| Number of frames per movie | 51 |  |  |  |  | 51 |  |  |  |
| Total dose (e-/Å<sup>2</sup>) | 50.5 |  |  |  |  | 50.5 |  |  |  |
| Defocus range (um) | 1.0 to 2.5 |  |  |  |  | 1.0 to 2.5 |  |  |  |
| **EM data processing** |  |  |  |  |  |  |  |  |  |
| Number of movies/micrographs | 6,015 |  |  |  |  | 12,002 |  |  |  |
| Box size (px) | 400 |  |  |  |  | 400 |  |  |  |
| Particle number (post 2D) | 823k |  |  |  |  | 424k |  |  |  |
| Particle number (post 3D) | 547k |  |  |  |  | 338k |  |  |  |
| Particle number (used in final map) | 39k | 38K | 32k | 24k | 43k | 38k | 43k | 31k | 64k |
| Symmetry | C1 | C1 | C1 | C1 | C1 | C1 | C1 | C1 | C1 |
| Map resolution (Å, FSC 0.143) | 2.58 | 2.58 | 2.68 | 3.1 | 2.65 | 2.77 | 2.73 | 2.71 | 2.56 |
| Local resolution range (Å, FSC 0.5) | 2.28-30 | 2.25-30 | 2.31-30 | 2.59-30 | 2.31-30 | 2.45-30 | 2.4-30 | 2.36-30 | 2.26-30 |
| Map sharpening B-factor (Å<sup>2</sup>) | -57.4 | -42.8 | -53.3 | -32.7 | -54.7 | -61 | -64.5 | -53 | -60.8 |
| **Model Building and Validation** |  |  |  |  |  |  |  |  |  |
| Initial model used | 8R1J |  |  |  |  |  |  | 8R1J |  |
| Model composition |  |  |  |  |  |  |  |  |  |
| Non-hydrogen protein atoms | 28937 |  |  |  |  |  |  | 33016 |  |
| Protein residues | 1790 |  |  |  |  |  |  | 2055 |  |
| RNA | 6 |  |  |  |  |  |  | 6 |  |
| RMSD from ideal |  |  |  |  |  |  |  |  |  |
| Bond length (Å) | 0.004 |  |  |  |  |  |  | 0.002 |  |
| Bond angles (°) | 0.533 |  |  |  |  |  |  | 0.499 |  |
| Validation |  |  |  |  |  |  |  |  |  |
| Molprobity score | 1.67 |  |  |  |  |  |  | 1.38 |  |
| Clashscore | 3.68 |  |  |  |  |  |  | 3.38 |  |
| Rotamers outliers (%) | 1.88 |  |  |  |  |  |  | 0.77 |  |
| FSC (0.5) model-vs-map | 2.76 |  |  |  |  |  |  | 2.88 |  |
| CC model-vs-map (masked) | 0.8 |  |  |  |  |  |  | 0.8 |  |
| Ramachandran plot |  |  |  |  |  |  |  |  |  |
| Favored (%) | 0.06 |  |  |  |  |  |  | 0.15 |  |
| Allowed (%) | 4.3 |  |  |  |  |  |  | 3.59 |  |
| Outliers (%) | 95.64 |  |  |  |  |  |  | 96.26 |  |

## Acknowledgments

We thank Adolfo García-Sastre for plasmids. We also thank George G. Brownlee and members of the Fodor and Grimes laboratories for helpful comments and discussions. This work was supported by Wellcome Trust Early Career Award 227570/Z/23/Z (KYC), Medical Research Council (MRC) programme grants MR/R009945/1 and MR/X008312/1 (EF), and Wellcome Investigator Awards 200835/Z/16/Z and 222510/Z/21/Z (JMG). Microscopy experiments were performed at the OPIC electron microscopy facility, an Instruct centre supported by the Wellcome Trust and the MRC. Computational analyses were carried out using the Oxford Biomedical Research Computing (BMRC) facility. Molecular graphics and analyses were performed using UCSF Chimera, developed by the Resource for Biocomputing, Visualization, and Informatics at the University of California, San Francisco, with support from NIH grant P41-GM103311. The views expressed are those of the authors and do not necessarily reflect those of the NHS, the NIHR, or the Department of Health.

## Author contributions

Conceptualization: KYC, AR, FW, ES and EF. Methodology: KYC, AR, LC. Investigation: KYC, AR, FW, LC. Visualization: KYC, AR. Resources: KYC and EF. Funding acquisition: KYC, JMG and EF. Data curation: KYC, AR. Validation: KYC, AR, FW, LC. Formal analysis: KYC, AR. Writing: KYC, AR, and EF. Supervision: KYC and EF. Project administration: KYC, JMG and EF.

## Competing interests

The authors declare no competing interests.

## Materials & Correspondence

Correspondence and requests for materials should be addressed to KYC, EF and JMG.

## Data availability

All data are available from the authors upon request.

## Notes

### Competing Interest Statement

The authors have declared no competing interest.

